# Hyperactive/impulsive and inattention symptoms are associated with reduced ERP activity during different reward processing stages: Evidence from the electrophysiological Monetary Incentive Delay Task in adult ADHD

**DOI:** 10.1101/817973

**Authors:** M. P. Bennett, H. Kiiski, Z. Cao, F. R. Farina, R. Knight, A. Sweeney, D. Roddy, C. Kelly, R. Whelan

## Abstract

Hyperactivity/impulsivity and inattention are core symptoms dimensions in attention-deficit/hyperactivity disorder. Some approaches suggest that these symptoms arise from deficits in the ability to anticipate and process rewards. However, evidence is equivocal with regard to ADHD-related differences in brain activity during reward processing. The aim of this study was to investigate when, and how, reward-related ERP activity was associated with hyperactive/impulsive symptoms and inattention symptoms. Adults with ADHD (n=34) and matched comparison participants (n=36) completed an electrophysiological version of the Monetary Incentive Delay task. This task separates reward processing into two stages-namely, an anticipation stage and a delivery stage. During the anticipation stage, visual cues signalled a possible monetary incentive (i.e. a reward or loss). After a brief delay, the delivery stage began, and incentives were delivered contingent on a speeded button-press. Electroencephalogram activity was simultaneously sampled and incentive-related event relate potentials (ERPs) calculated. These data were then analysed by calculating multiple regression models, at each sample point, wherein the correlation between incentive-related ERPs and ADHD symptoms was estimated. Linear and curvilinear associations between ERP activity and ADHD symptoms were tested in each regression mode. Findings suggest that ADHD symptoms were associated ERP activity at different reward processing stages. Hyperactive/impulsive symptoms were associated with reduced ERP activity during the initial anticipation of rewards from 224-329 ms post-reward cue. Inattention symptoms were associated with reduced ERP activity during the initial delivery of rewards from 251-280 ms post-reward onset. Finally, extreme ends of hyperactive/impulsive and inattention symptoms were associated with reduced ERP activity towards the end of the anticipation stage from 500 ms post-reward cue onwards. These results support the idea that reward processing is disrupted in ADHD while also shedding new light on the dynamic relationship between ADHD symptom dimensions and the neurological mechanisms of reward processing.

---

Attention-Deficit/Hyperactivity Disorder (ADHD) is a neurodevelopmental disorder characterized by patterns of impulsivity, hyperactivity and/or inattention (1). Although ADHD is typically diagnosed in childhood, behavioural symptoms continue into adulthood for most individuals, with an overall prevalence rate of 3-4% (2–5). Recent etiological models posit that ADHD symptoms result from a diminished ability to anticipate rewards (6–8). According to these approaches, ADHD symptoms accrue from lower phasic dopaminergic activity in response to reward-predictive cues (9). This view is supported by evidence of ventral striatum (VS) hypoactivity in response to cues that signal monetary rewards in both adult and adolescent ADHD, relative to healthy comparisons (10–13). Despite these findings, however, the relationship between reward processing and ADHD remains unclear. Furthermore, it is unknown whether deficits in reward processing mediate specific symptoms dimensions in ADHD (e.g. impulsivity and/or inattention) or the disorder in general.

Conflicting evidence regarding the relationship between reward processing and ADHD symptoms has been reported using functional magnetic resonance imaging (fMRI) and the monetary incentive delay (MID) task (14). The MID task separates reward processing into two discrete stages. Across trials, a visual cue first signals a possible monetary incentive: the anticipation stage. After a brief period, the incentive is then delivered on a partial reinforcement-schedule, contingent on a response to a target: the delivery stage. A number of studies report a negative association between VS activity in response to cues that predict monetary rewards and hyperactive/impulsive symptoms – but not inattentive symptoms (10–13). These results imply that inefficient reward anticipation underlies this specific ADHD symptom dimension. However, the opposite pattern has also been found. In a recent study with a large sample of adolescents, the authors reported a negative association between striatal activity in response to reward-predictive cues and inattentive symptoms – but not hyperactive/impulsive symptoms (13). Moreover, others report no evidence of reduced brain activity in response to reward predictive-cues in ADHD. These studies instead highlight either VS hyperactivity in response to reward-predictive cues in ADHD relative to comparisons or no between-group differences (15–18).

The lack of consensus regarding ADHD-related deficits during reward processing may partly stem limits inherent in previous FMRI research. While spatially informative, fMRI lacks the temporal resolution to study the transient neuro-cognitive process that mediate each stage of the MID task. Indeed, the stages of reward processing (such as anticipation and delivery) can be further divided into brief cognitive mechanisms like cue detection, attentional allocation, motor-response preparation and evaluative appraisal (19). It is possible that ADHD symptoms relate to atypical patterns of activity during these discrete neuro-cognitive mechanisms involved in reward processing. Therefore, alternative approaches that afford greater temporal granularity are warranted.

An electrophysiological (EEG) version of the MID task (e-MID) represents one such promising approach. This task uses event-related potentials (ERPs; time-locked electroencephalograms) to precisely quantify the temporal dynamics of reward processing (15, 19–22). Two ERP components are particularly relevant during the anticipation stage: cue-P3 and contingent negative variation (CNV). Cue-P3 amplitudes occur in response to predictive cues, with a peak near 300 ms that is maximal at posterior scalp spaces. Cue-P3 amplitudes are thought to reflect attentional allocation (19–21). CNV ERPs are negative-going potential shifts observed 1000-2000 ms after the cue-P3 and are maximal at frontal scalp spaces. These amplitudes are indicative of motor-response preparation. Importantly, the amplitudes of both cue-P3 and CNV are increased by the motivational magnitude of the cue (20, 23). Three additional ERP components are relevant to the delivery stage: feedback-related negativity (FRN), reward-positivity (RewP) and feedback-P3. FRN amplitudes are observed from 200-300 ms after feedback and are maximal in frontro-central scalp spaces. These amplitudes are thought to reflect a evaluative processing outcome; that is, the negativity of FRN amplitudes increases as a function of the unfavorability of an outcome (21). The RewP component occurs in approximately the same period as the FRN, albeit in the context of reward delivery; the RewP component is characterised by an emergent positivity that increases with relative gains (19, 24, 25). P3 amplitudes following incentive feedback (FB-P3) have also been reported, reflecting enhanced attentional allocation towards salient outcomes (20, 21). Using these incentive-related ERPs as guide, the current study sought to better characterise the relationship between ADHD symptoms and neuro-cognitive mechanisms of reward processing.

Another important consideration is the possible curvilinear association between ADHD symptoms and activity in reward-related brain areas. While there is evidence of VS hypoactivity in individuals diagnosed with ADHD relative to comparison groups (10–12), the opposite finding exists in studies with community samples. These latter studies instead report a relative VS hyperactivity in individuals with elevated ADHD symptoms (26, 27). It has therefore been suggested that the relationship between ADHD symptoms and brain activity is dimensional and follows an inverted-u-shape (9). That is, maximal brain activation is present at mid-range ADHD symptom-severity, with reduced brain activation present at the extreme ends of ADHD symptom-severity scale (9). However, there is a lack of direct evidence for this inverted-u-shaped association. One reason is that between-group designs, which compare high and low symptom reporters, are insensitive to curvilinear associations between brain activity and symptoms. A continuous analysis is required to better examine the dimensional association between brain activity and ADHD symptoms.

No previous study has examined reward processing in adult ADHD using the e-MID task. We therefore administered this task to a sample of adults with a wide range of ADHD symptoms. We aimed to determine specific time points when ERP activity was associated with hyperactive/impulsive and inattention symptoms that are characteristic of with ADHD. We analysed the data continuously using regression and tested for both linear and curvilinear associations between ADHD symptoms and ERP activity. Findings revealed that hyperactive/impulsive and inattention symptoms were associated with reduced ERP activity during different stages of reward processing.

## Methods

### Participants

Participants were recruited through a targeted advertising campaign for people self-identifying as either having or not having a diagnosis of ADHD. Exclusion criteria included epilepsy, severe migraine, severe motor impairments, stoke, neuro-developmental disorders other than ADHD and history of concussion and/or head trauma resulting in loss of consciousness. Participants were also excluded if prescribed selective serotonin re-uptake inhibitors, benzodiazepines, anti-psychotics, anti-convulsants and/or atomoxetine. These criteria were determined via an initial telephone interview. Potential participants also completed an abbreviated version of the Structured Clinical Interview for the DSM-IV, which was used to check for symptoms of psychosis, mood disorder, dysthymia, bipolar disorder, suicidality, eating disorders, substance use, alcohol use, anxiety disorders and obsessive-compulsive disorder. Participants were excluded on the basis of psychosis and/or bipolar disorder, although no participants met these exclusion criteria.

Thirty-four participants with a previous diagnosis of ADHD were recruited as were thirty-six participants with no previous diagnosis of ADHD. Participants were well-matched in terms of age, sex and years of education (Table 1). ADHD symptom severity at the time of testing was established using the long form of the self-reported Connors Adult ADHD Rating Scale (CAARS). Based on the CAARS DSM symptom subscale of the CAARS, a t-score of ≥ 65 is recommended as a threshold for clinically significant symptoms. Using this criterion, eight participants without a diagnosis of ADHD reported clinically significant symptoms at the time of testing. Conversely, six participants with a diagnosis of ADHD did not to reach the clinical threshold for ADHD at the time of testing (see Figure 1). Hyperactive/impulsive symptoms and inattentive symptoms were also indexed using the DSM subscales provided by the long form CAARS (Figure 1; Table 1).

**Table 1.**
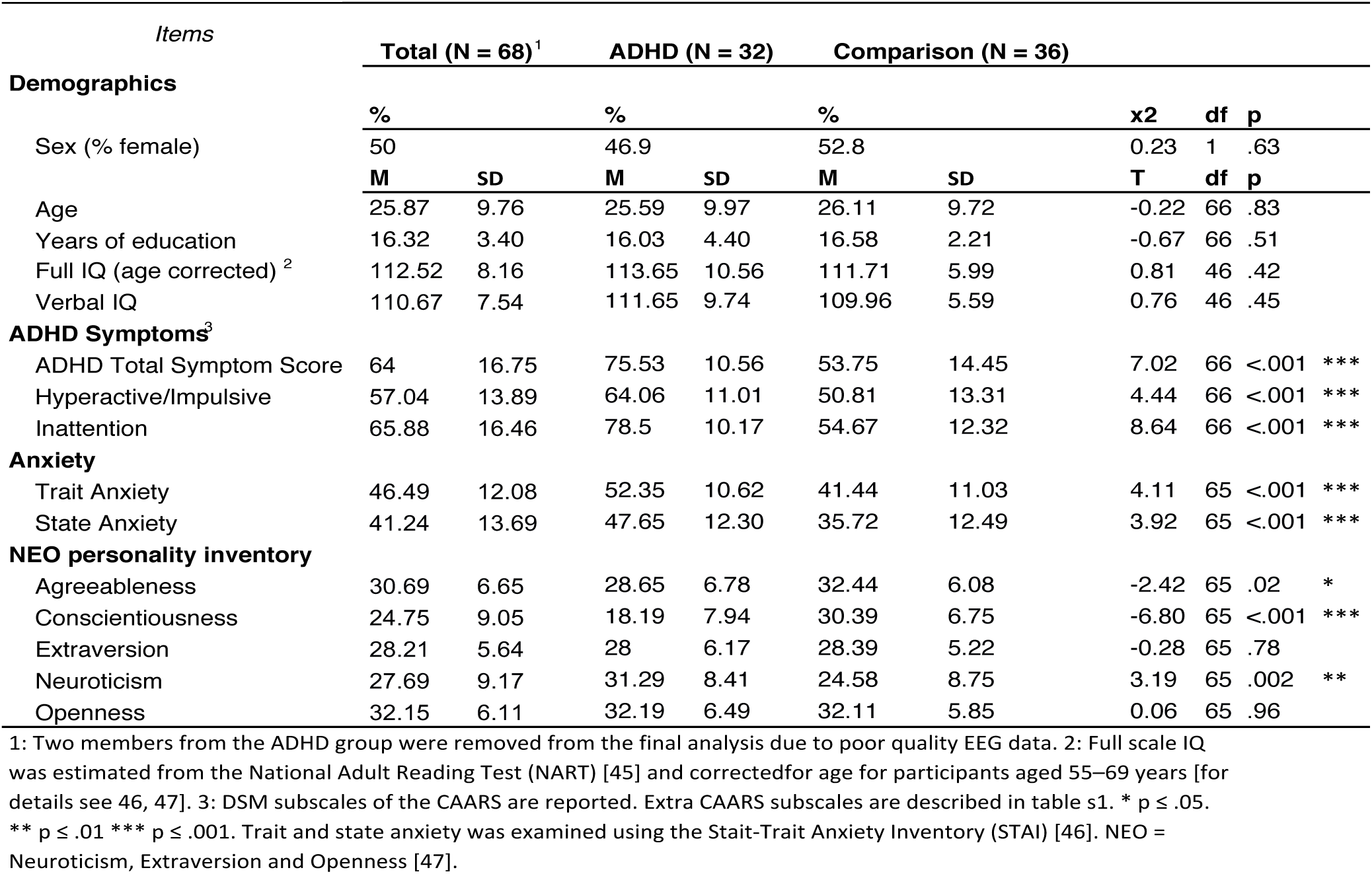
Demographic details

| Items | Total (N = 68)<sup>1</sup> |  | ADHD (N = 32) |  | Comparison (N = 36) |  |  |  |  |
| --- | --- | --- | --- | --- | --- | --- | --- | --- | --- |
| Demographics |  |  |  |  |  |  | x2 | df | p |
| Sex (% female) | 50 |  | 46.9 |  | 52.8 |  | 0.23 | 1 | .63 |
|  | M | SD | M | SD | M | SD | T | df | p |
| Age | 25.87 | 9.76 | 25.59 | 9.97 | 26.11 | 9.72 | -0.22 | 66 | .83 |
| Years of education | 16.32 | 3.40 | 16.03 | 4.40 | 16.58 | 2.21 | -0.67 | 66 | .51 |
| Full IQ (age corrected)<sup>2</sup> | 112.52 | 8.16 | 113.65 | 10.56 | 111.71 | 5.99 | 0.81 | 46 | .42 |
| Verbal IQ | 110.67 | 7.54 | 111.65 | 9.74 | 109.96 | 5.59 | 0.76 | 46 | .45 |
| **ADHD Symptoms<sup>3</sup>** |  |  |  |  |  |  |  |  |  |
| ADHD Total Symptom Score | 64 | 16.75 | 75.53 | 10.56 | 53.75 | 14.45 | 7.02 | 66 | <.001 *** |
| Hyperactive/Impulsive | 57.04 | 13.89 | 64.06 | 11.01 | 50.81 | 13.31 | 4.44 | 66 | <.001 *** |
| Inattention | 65.88 | 16.46 | 78.5 | 10.17 | 54.67 | 12.32 | 8.64 | 66 | <.001 *** |
| **Anxiety** |  |  |  |  |  |  |  |  |  |
| Trait Anxiety | 46.49 | 12.08 | 52.35 | 10.62 | 41.44 | 11.03 | 4.11 | 65 | <.001 *** |
| State Anxiety | 41.24 | 13.69 | 47.65 | 12.30 | 35.72 | 12.49 | 3.92 | 65 | <.001 *** |
| **NEO personality inventory** |  |  |  |  |  |  |  |  |  |
| Agreeableness | 30.69 | 6.65 | 28.65 | 6.78 | 32.44 | 6.08 | -2.42 | 65 | .02 * |
| Conscientiousness | 24.75 | 9.05 | 18.19 | 7.94 | 30.39 | 6.75 | -6.80 | 65 | <.001 *** |
| Extraversion | 28.21 | 5.64 | 28 | 6.17 | 28.39 | 5.22 | -0.28 | 65 | .78 |
| Neuroticism | 27.69 | 9.17 | 31.29 | 8.41 | 24.58 | 8.75 | 3.19 | 65 | .002 ** |
| Openness | 32.15 | 6.11 | 32.19 | 6.49 | 32.11 | 5.85 | 0.06 | 65 | .96 |

1: Two members from the ADHD group were removed from the final analysis due to poor quality EEG data. 2: Full scale IQ was estimated from the National Adult Reading Test (NART) [45] and corrected for age for participants aged 55–69 years [for details see 46, 47]. 3: DSM subscales of the CAARS are reported. Extra CAARS subscales are described in table s1. \* $p \leq .05$ .
\*\* $p \leq .01$ \*\*\* $p \leq .001$ . Trait and state anxiety was examined using the State-Trait Anxiety Inventory (STAI) [46]. NEO = Neuroticism, Extraversion and Openness [47].

**Figure 1.**
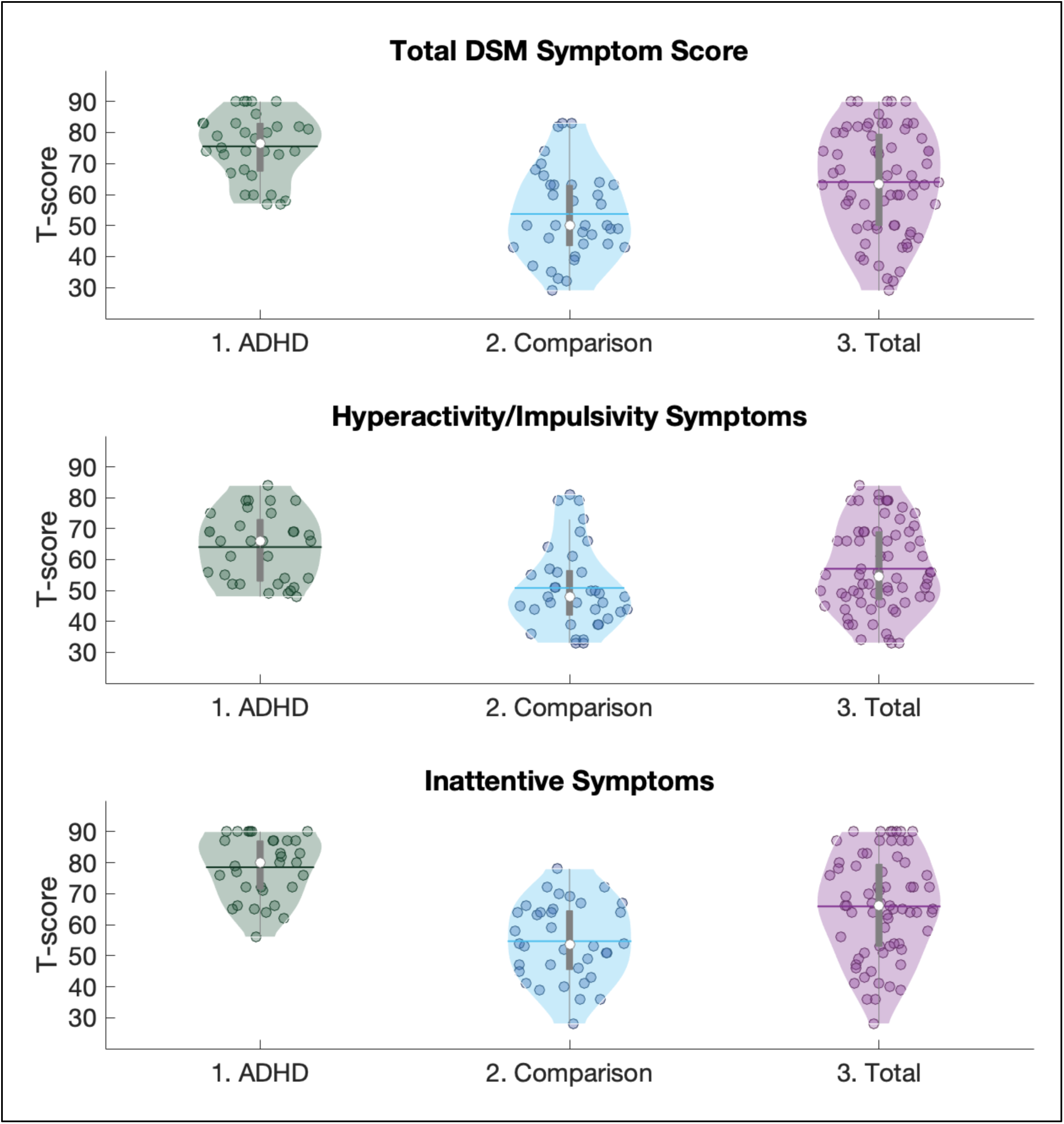
Distribution of ADHD symptoms within the entire participant sample. Participants either (1) had a diagnosis of ADHD or (2) did not have a diagnosis of ADHD. Participants were then (3) pooled together into one overall cohort. This approach ensured a good spread of ADHD symptoms. Symptom scores are based on Conner’s adult ADHD rating scale DSM criteria. The white dot at the centre of each violin plot indicates the mean. The straight line at the centre of each violin plot indicates the median. Boxes indicate the interquartile range and error bars indicate 95% confidence intervals.

Demographic and diagnostic details are reported in Table 1 (also Table S1). Ethical approval was obtained from the School of Psychology Research Ethics Committee at Trinity College Dublin. All participants provided written consent and received €40 (approx. $50) reimbursement. Where relevant, participants abstained from ADHD-relevant medications for 36 hours prior to data collection. At the time of testing, 11 individuals with a diagnosis of ADHD were taking methylphenidate and one was taking lisdexamfetamine. In addition, two other individuals with ADHD reported taking methylphenidate in the past while one took norepinephrine re-uptake inhibitors and another took serotonin re-uptake inhibitors in the past.

### E-MID Task

The full testing protocol is described in the Supplementary Materials. Participants sat in a dark, sound-attenuated room, 1.05 m from a CRT monitor (75 Hz refresh rate; visual angle of 5.4° horizontally and 5.6° vertically). Stimuli appeared in the middle of the screen.

At the beginning of each e-MID trial, a coloured square was presented for 250 ms (Figure 2). A reward cue (green square) signalled the potential to win 20 c. A loss cue (red square) signalled the potential to lose 20 c. A neutral cue (blue square) signalled that neither a reward nor loss was possible. This established an incentive anticipation stage. A blank inter-stimulus interval (ISI) was then displayed for 2000-2500 ms (randomly, using a uniform distribution), followed by a target (white star) for 160-250 ms (the response interval). Participants were instructed to respond to the target as quickly as possible with their left or right index finger via a response pad. An adaptive algorithm tracked response accuracy, trial-by-trial, to achieve a 66% success rate. The response interval was shortened if the success rate exceeded 66% (making hits more difficult) and lengthened if the success rate was below 66% (making hits more achievable) (see Supplemental Materials; Table S2). Feedback was presented for 1500 ms after target offset (i.e., the incentive delivery stage). Response hits on reward trials produced a text reading “20 c” in green font (i.e., reward delivery). Response misses on reward trials produced a sad emoticon (i.e., reward-omission: ~33% of trials). Response misses on loss trials produced a message reading ‘-20 c’ in red front (i.e., loss delivery: ~33% of trials). Response hits on the loss trials produced a happy emoticon (i.e., loss-omission). Hits and misses on the neutral trials produces a happy and sad emoticon, respectively (i.e., non-monetary outcomes). Here, participants were told that although they would receive feedback about the speed of their response, they could neither win nor lose money. Thirty practice trials were completed (10 trials per condition), followed by 48 trials of each incentive condition. Trials were separated by a 2000 ms inter-trial interval.

**Figure 2.**
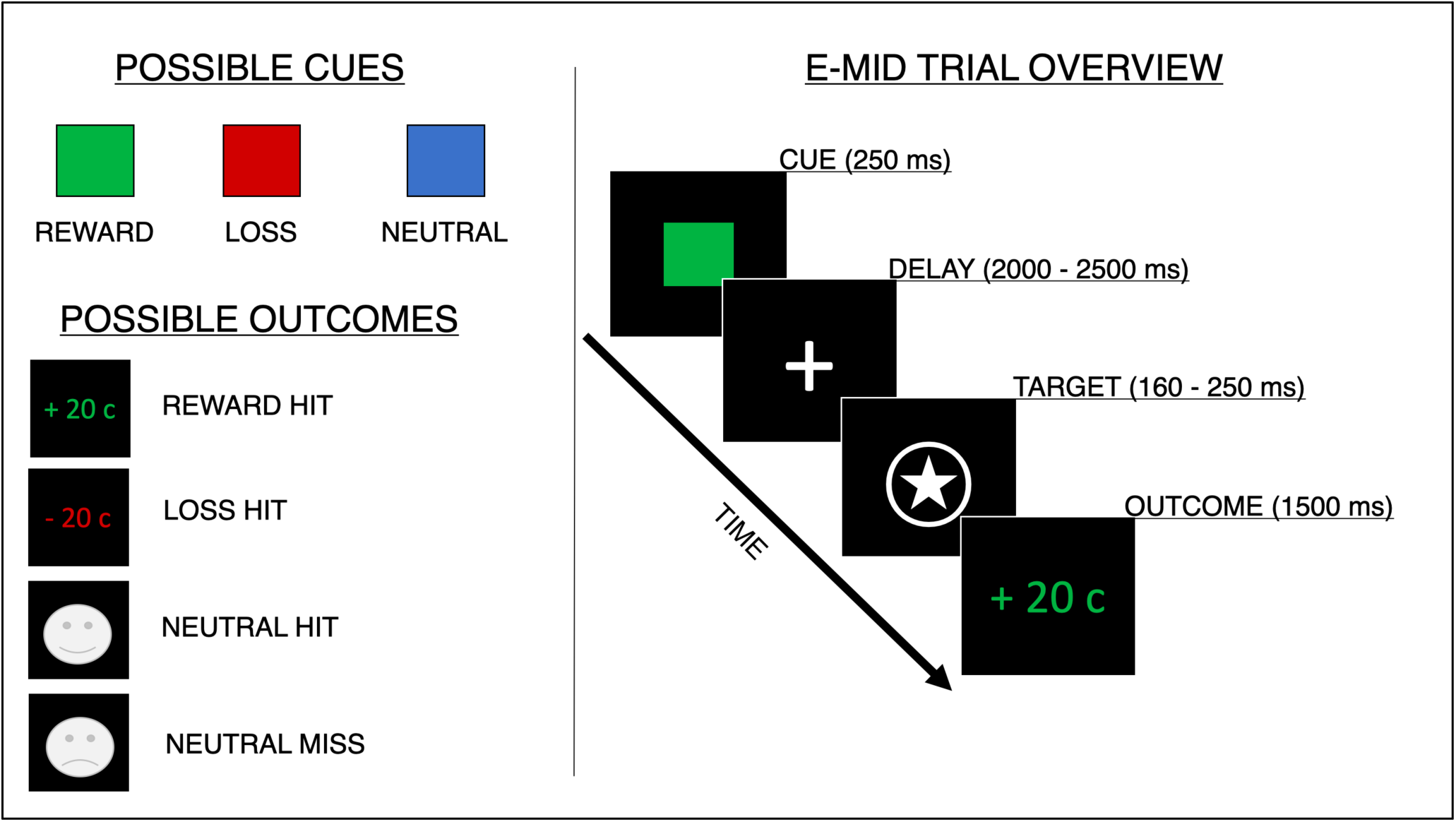
Schematic overview of the e-MI

### Data analysis

#### Reaction times

Mean reaction times (RTs) to target were calculated for reward, loss and neutral trials. Repeated measures analysis of variance (rmANOVA) examined the effect of incentive (reward, loss and neutral) on mean RT. Pearson’s correlations were used to estimate the association between incentive-related RTs and hyperactive/impulsive symptoms as well as inattention symptoms.

### EEG

#### Acquisition and Pre-processing

EEG was recorded using an ActiveTwo Biosemi™ system. Sixty-four (Ag/AgCl) scalp electrodes were placed using a fitted electrode cap (Biosemi™) according to the 10–20 system (28). Two additional electrodes were placed on the mastoids. Vertical and horizontal electro-oculograms were recorded bilaterally (approximately 2 cm below each eye and from each outer canthus). Pre-processing used the EEGLAB toolbox (29) (http://sccn.ucsd.edu/eeglab) and the FASTER plug-in (30) (http://sourceforge.net/projects/faster) (see Supplementary Materials for details on pre-processing and data rejection).

The pre-processed EEG data for each participant were partitioned into two epochs: from −200 ms to 2000 ms around cue onset (anticipation stage) and −200 ms to 1500 ms around feedback onset (delivery stage). Epochs were extracted for each of the following conditions: (a) reward cue, (b) loss cue, (c) neutral cue, (d) reward hit, (e) reward miss, (f) loss hit, (g) loss miss, (h) neutral hit and (i) neutral miss. Each epoch was baseline adjusted by subtracting the mean amplitude of the 20-ms pre-stimulus period. Data were referenced to the average of the mastoids. For each participant, an average epoch was created for each condition by calculating the mean value across the total number of available epochs. This step was repeated at each electrode channel.

Averaged epochs were grouped together into scalp-based regions of interest (ROIs). This was accomplished by calculating the mean across clusters of electrode channels (31): namely, fronto-polar (FPz, FP1, FP2, AF3, AF4), left fronto-lateral (AF7, F5, F7), right fronto-lateral (AF8, F6, F8), fronto-central (FCz, Cz, FC1, FC2, C1, C2), occipital-medial (Oz, POz, O1, O2, PO3, PO4), left posterior-lateral (CP5, CP9, P5, P7, P9) and right posterior-lateral (CP8, CP10, P6, P8, P10). Cue-P3 and FB-P3 amplitudes are maximal at posterior electrode channels (15, 20–22); therefore, three posterior-scalp ROIs were generated (i.e., occipital-medial, left and right posterior-lateral). CNV and FRN amplitudes are maximal at fronto-central electrode channels; therefore, four frontal-scalp ROIs were generated (i.e., fronto-polar, fronto-central, left and right fronto-lateral) (15, 20–22). Finally, a 20 Hz high pass filter was applied to each signal.

#### Analysis strategy

For each participant, anticipation stage ERP epochs consisted of 1126 sample points (i.e. −200 ms until 2000 ms) for each incentive condition (i.e. reward-cue, loss-cue and neutral-cue). Delivery stage ERP epochs consisted of 870 sample points (i.e. −200 until 1500 ms) for each incentive condition (i.e. reward-hit, loss-hit and neutral-hit). These data were analysed using an approach that isolated when incentive-related ERP activity was associated with ADHD symptoms. At each time (or sample) point, a regression model was calculated wherein the correlation between ERP activity and ADHD symptoms was estimated. Hyperactive/impulsive or inattentive symptoms were separately examined. Target (symptom scores) and predictor (ERP activity) variables were all transformed to z-scores.

A linear and curvilinear association between the predictors and ADHD symptoms was tested in each regression model (Figure 3). The u-shaped association was estimated by squaring the ERP z-scored values. As a consequence, positive beta values indicate a typical-u-shaped association between ERP activity and symptoms scores. Conversely, negative beta values indicate an inverted-u-shaped association between ERP activity and symptoms scores (Figure 3). Sex, age and years of education were added as nuisance covariates. Also, we included ERP activity during the neutral condition to control for any fundamental association between ADHD and e-MID task performance. Covariates were also transformed to z-scores. Thus, the model calculated at each time point was as follows:

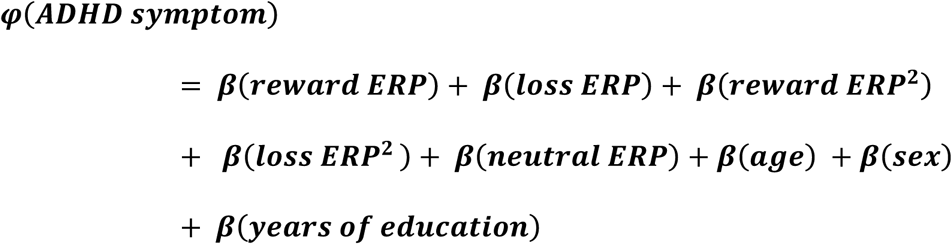

Here, ‘*ADHD symptom’* refers to either hyperactive/impulsive or inattentive symptom scores based on the DSM subscales of the long-form CAARS. In the context of the anticipation stage, ‘*reward ERP’*, ‘*loss ERP’* and ‘*neutral ERP’* refers to ERP activity in response to the reward, loss and neutral-cues. In the context of the delivery stage, ‘*reward ERP’*, ‘*loss ERP’* and ‘*neutral ERP’* refers to ERP activity in response to the delivery of reward, loss and neutral outcomes (or ‘hits’). Models were calculated at each individual sample point (i.e. 1126 samples during the anticipation stage and 861 samples during the delivery stage) and at each of the seven scalp ROI.

**Figure 3.**
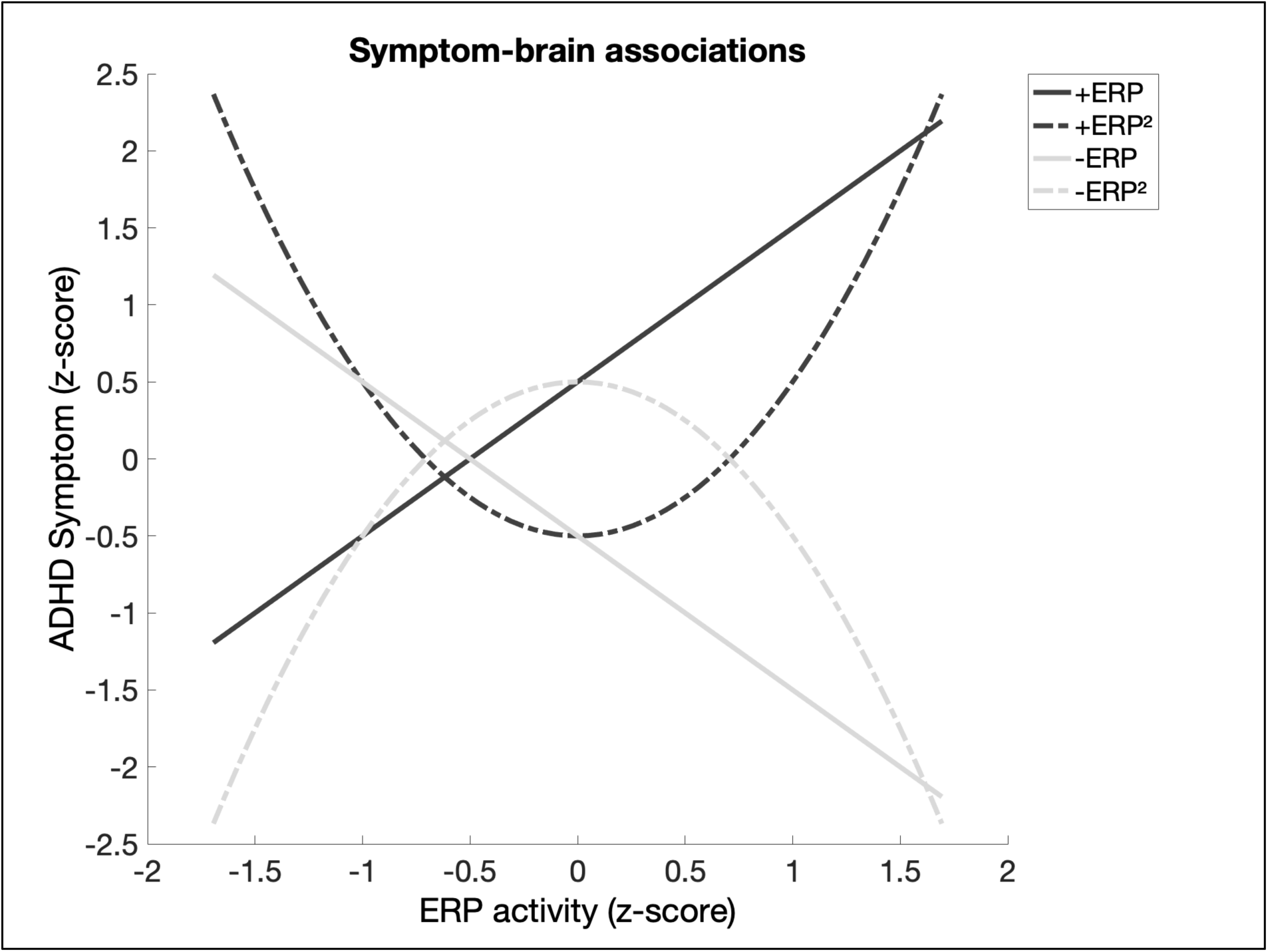
Possible associations between ADHD symptoms and incentive-related EEG activity based on simulated data. Within each regression model, terms were included to estimate a linear (solid lines; ERP) or curvilinear (dashed lines; ERP^2^) association. A positive beta value indicates that the linear association was positive going (black-solid lines) or that the curvilinear association was typical-u-shaped (black-dashed lines). A negative beta value indicates that the linear association was negative going (grey-solid lines) or that the curvilinear association was inverted-u-shaped (grey-dashed lines).

#### Statistical significance

The statistical significance of the regression model terms was determined using two methods. First, we used a maximum statistic approach that involved the generation of random permutation models. That is, for every ROI, 1000 null models were also calculated by randomly shuffling the EEG predictors across participants before re-calculating the regression model. This created a set of 7000 null models at each time point (1,000 null models x 7 ROIs). Pooling all 7000 null models together allowed us to correct for multiple comparisons across ROIs by identifying values that exceeded the bottom 2.5 or top 97.5 percentile of null distribution. This method was applied to (1) identify ROIs in which the R squared (R^2^) value was significantly greater than chance and (2) identify significant beta value within each model.

The second approach used to determine statistical significance was based the number of contiguous time points where a model term exceeded the maximum statistical significance threshold. If the maximum statistic approach revealed a significant finding (e.g. R^2^ term or beta value that passed its threshold), then the number of contiguous significant time points (*x*) was counted. This total was described in terms of the probability of it appearing in all 7000 null models. That is, P(number of null models featuring *x* contiguous samples | number of null models). We referred to this as ‘*contiguous* p’.

Statistical tests were carried out separately for the anticipation and delivery stages of ERP activity. Tests were calculated using MATLAB (Version 9.1, Natick, Massachusetts, US: The MathWorks Inc.). Data and processing scripts are available online (https://osf.io/k4hqz/).

## Results

### Reaction times

**Table 2.**
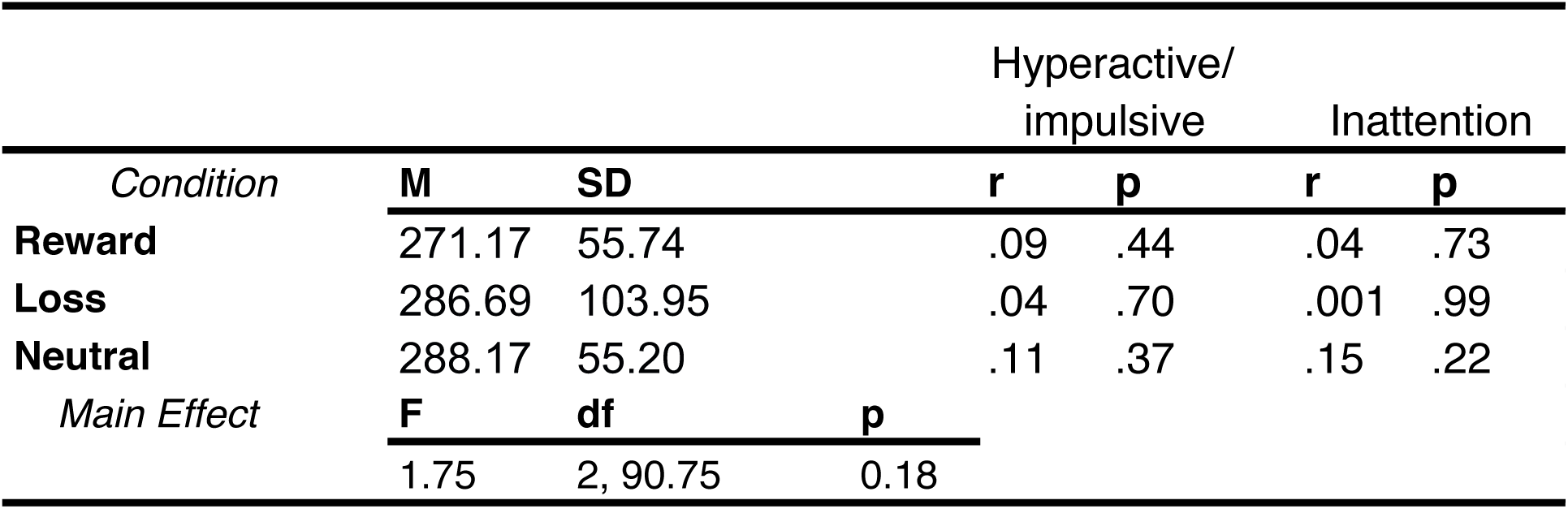
Reaction times (in ms) during anticipation stage

### EEG

For two participants, more than 50% of trials were rejected due to poor quality. These participants were excluded from further analysis (see Supplemental Materials). The R^2^ values and beta values for each EEG-related term at each ROI are presented in Supplementary Materials. Only the significant effects are described below. Figures 4 and 7 illustrate the averaged ERP activity for the anticipation stage (impulsive/hyperactivity and inattentive symptoms, respectively). Figures 6 and 9 illustrate the averaged ERP activity for the delivery stage (impulsive/hyperactivity and inattentive symptoms, respectively). Detailed results from each scalp ROI are described in Supplementary Materials. This includes the association between hyperactive/impulsive symptoms and ERP activity during (i) the anticipation (Figures S1-S7) and (ii) delivery stage (Figure S8-S14). Also included is the association between inattention symptoms and ERP activity during (iii) the anticipation stage (Figures S15-S21) and (iv) delivery stage (Figure S22-S28). Below, we discuss the findings from scalp ROIs where the R^2^ value surpassed the critical R^2^ threshold.

**Figure 4.**
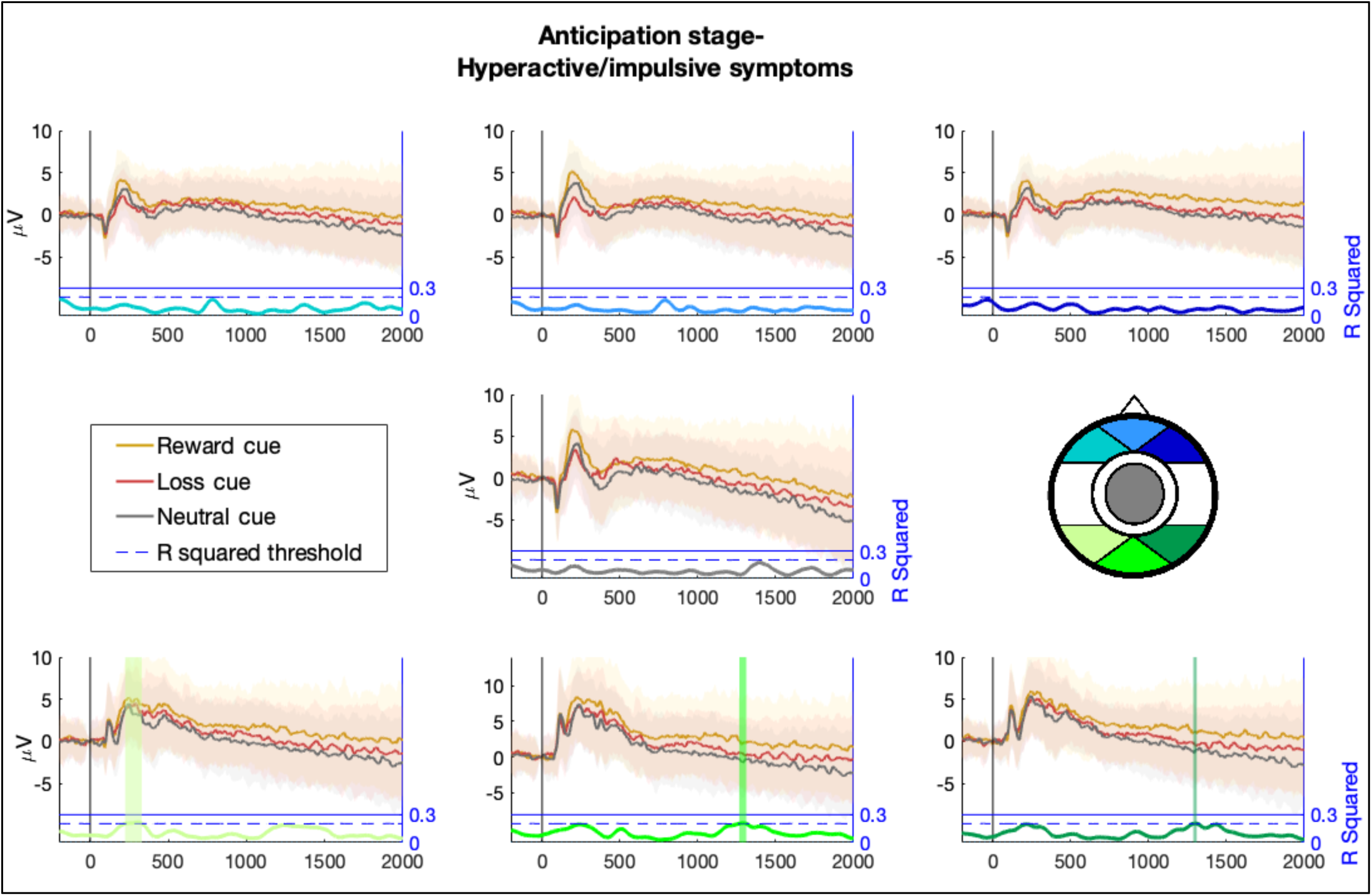
Average ERP amplitudes in response to reward, loss and neutral cues during the anticipation stage. Each panel represents a scalp ROI. Lighter region surrounding the ERP amplitudes represents the standard deviation. μV = micro volts. The coloured vertical bars illustrate contiguous periods when ERP amplitudes were significantly associated with Hyperactive/impulsive symptoms. ERP activity at three scalp ROIs was significantly associated with hyperactive/impulsive symptoms. This is based on whether R^2^ values continuously exceeded the top 97.5 percentile of all (7000) null model R^2^ values (i.e R^2^ threshold). The R^2^

### Hyperactive/impulsive

#### Anticipation Stage

Based on the maximum statistic approach, the critical R^2^ threshold over the entire epoch was around 0.20 ± 0.005 (M ± SD). In addition, the probability of observing two (or more) consecutives samples with a significant R^2^ value was less than one in every 7000 models. Using these criteria, ERP activity from 3 (of scalp ROIs was significantly associated with hyperactive/impulsive symptoms (Figure 4). These were the left posterior lateral, occipital-medial and right posterior lateral occipital-medial and right posterior lateral ROIs (Figure 5).

**Figure 5.**
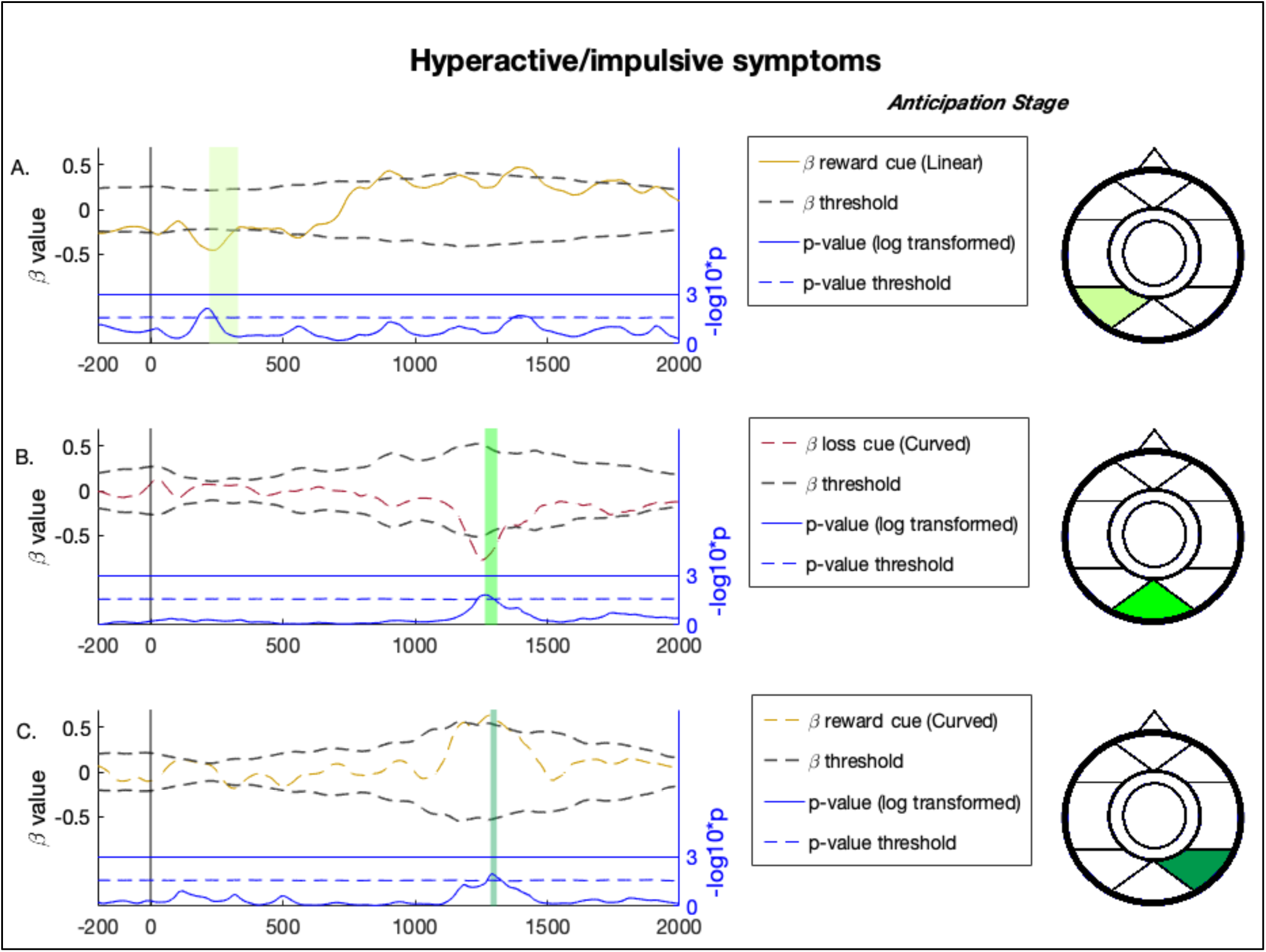
Hyperactive/impulsive symptoms were significantly associated with incentive-related ERP activity at 3 scalp ROIs during the anticipation stage (left hand side). Significant periods are illustrated by coloured vertical bars in each panel; these coloured bars correspond to the contiguously significant R^2^ values (also in Figure 4). Further analysis revealed that only certain incentive conditions were associated with hyperactivity/impulsivity symptoms at each scalp ROI. (A) At the posterior-lateral left ROI, the association between ERP activity and symptoms was driven by the reward-cue (linear association with symptoms). (B) At the occipital-medial ROI, the association between ERP activity and symptoms was driven by the loss-cue (curved association). (C) At the posterior-lateral right ROI, the association between ERP activity and symptoms was driven by the reward-cue (curved association). Statistical significance was based on the contiguous beta values (coloured dotted lines) that exceeded the bottom 2.5 or top 97.5 percentile of the corresponding beta values from all (7000) null models (i.e. the 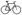threshold; the grey dotted line). Also illustrated are the p-values that corresponds to the beta values. These are presented in blue at the bottom of each panel. We applied the maximum statistic approach to these values to identify p-values that exceeded the top 97.5 percentile of the corresponding p values from all null models (i.e. a p threshold; the blue dotted line). This value was log transformed so that higher values trended towards statistical significance: where p = .05, −log10(p) = 1.30; where p = .01, −log10(p) = 2; where p = .001, − log10(p) = 3.

The left posterior lateral ROI passed the significance threshold from 224 until 329 ms (R^2^ range = 0.20 to 0.21) (Figure 4; Figure S5), a period corresponding to the cue-P3 component. Within this ROI, there was a significant negative association between reward-cue ERP activity and hyperactive/impulsive symptoms (β range = − 0.45 to −0.37; β threshold (M ±SD) = −.21 ± 0.001) over 35 contiguous samples (179 − 245 ms, *contiguous* p < .001) (Figure 5A; Figure S5). The negative association between ERP activity and hyperactive/impulsive symptoms was linear.

The occipital-medial ROI exceeded the significance threshold from 1267 until 1310 ms (R^2^ range = 0.20 to 0.21) (Figure 4; Figure S6), a period corresponding to an early CNV component. Within this ROI, a significant curvilinear association between loss-cue ERP activity and hyperactive/impulsive was found (β range = −0.77 to −0.65; β threshold (M ±SD) = −.49 ± 0.02) over 32 samples (1238 − 1297 ms, *contiguous* p < .001) (Figure 5B; Figure S6). This association was an inverted-u-shaped, indicated by the negative beta value: Loss-cue related ERP activity increased as symptoms transitioned from low to moderate but decreased as symptoms transitioned from moderate to high.

The right posterior lateral ROI surpassed the significance threshold from 1288 until 1308 ms (R^2^ range = 0.20 to 0.21) (Figure 4; Figure S7). This period corresponds to an early CNV component. Within this ROI, a significant curvilinear association between reward-cue ERP activity and hyperactive/impulsive was found (β range = 0.54 to 0.63; β threshold (M ±SD) = −.53 ± 0.01) over 28 samples (1275 − 1327 ms, *contiguous* p < .001) (Figure 5C; Figure S7). This association was typical-u-shaped, indicated by the positive beta value. Reward-cue ERP activity decreased as symptoms Transitioned from low to moderate, but increased as symptoms transitioned from moderate to high.

#### Delivery stage

Based on the maximum statistic approach, the critical R^2^ threshold over the entire epoch was around 0.20 ± 0.007 (M ± SD). The probability of observing two (or more) consecutives samples with a significant R^2^ value was less than one in every 7000 models. Using these criteria, delivery stage incentive-related ERP activity was never significantly associated with impulsivity at any ROI (Figure 6).

**Figure 6.**
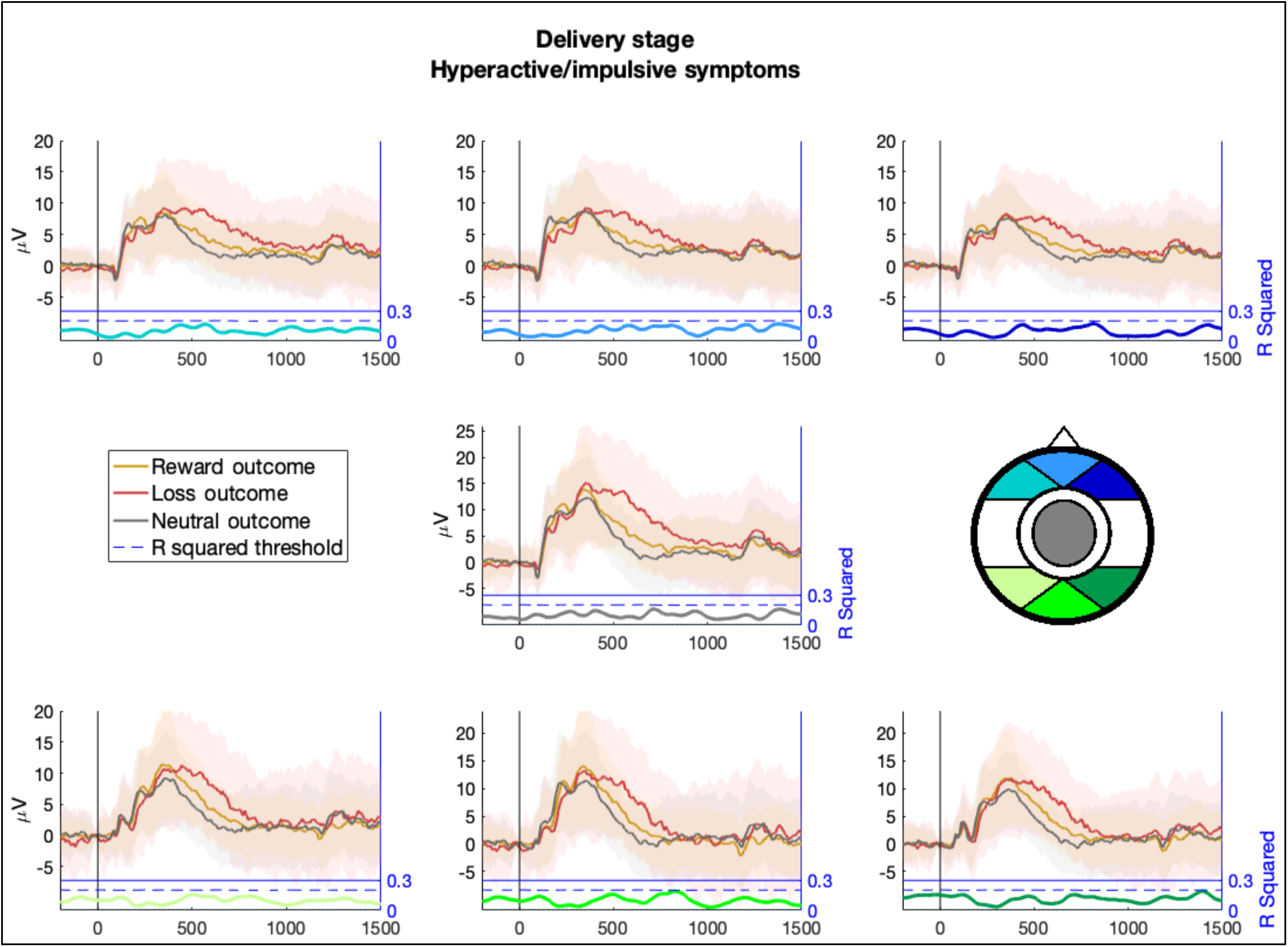
Average ERP amplitudes in response to reward, loss and neutral outcomes during the delivery stage. Each panel represents a scalp ROI. Lighter region surrounding the ERP amplitudes represents the standard deviation. μV = micro volts. ERP activity during the delivery stage was never significantly associated with hyperactive/impulsive symptoms. This is based on R^2^ values surpassing the significance threshold. The R^2^ value at each time point is illustrated at the bottom of each ROI panel- the colour corresponds to the scalp ROI.

### Inattentive Symptoms

#### Anticipation Stage

Based on the maximum statistic approach, the critical R^2^ threshold across the entire epoch was around 0.20 ± 0.007 (M ± SD). The probability of observing two (or more) consecutives samples with a significant R^2^ value was less than one in every 7000 models. Based on these criteria, ERP activity from two scalp ROIs was significantly associated with inattentive symptoms (Figure 7). In temporal order, these were the left fronto-lateral and left posterior-lateral ROIs.

**Figure 7.**
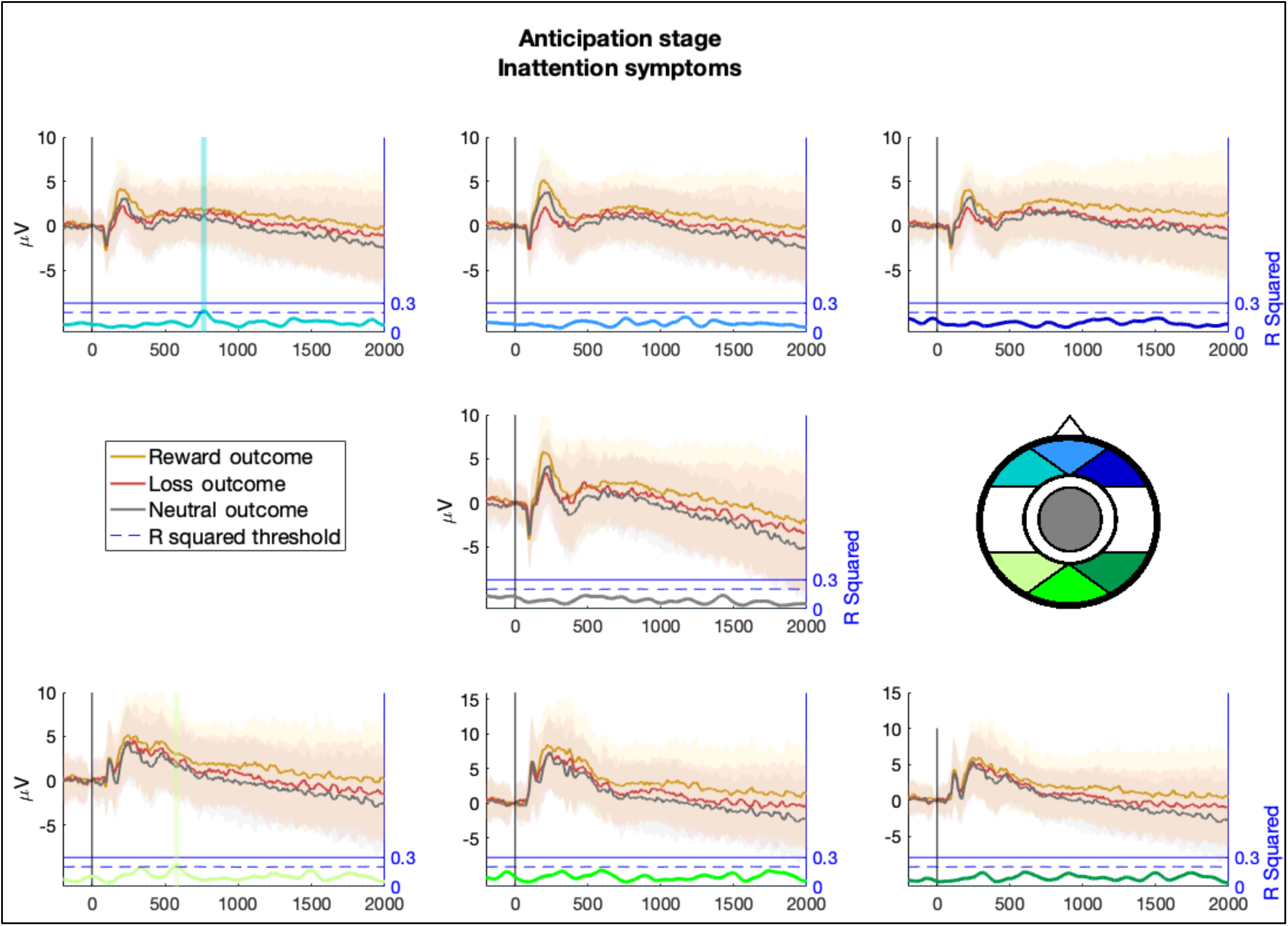
Average ERP amplitudes in response to reward, loss and neutral outcomes during the anticipation stage. Each panel represents a scalp ROI. Lighter region surrounding the ERP amplitudes represents the standard deviation. μV = micro volts. The coloured vertical bars illustrate contiguous periods when ERP amplitudes were significantly associated with inattention symptoms. Incentive-related ERP activity at the frontolateral left and posterior-lateral left ROIs was significantly associated with inattention symptoms. This is based on whether R^2^ values continuously exceeded the top 97.5 percentile of all (7000) null model R^2^ values (i.e. the R^2^ threshold). The R^2^ value at each time point is illustrated at the bottom of each ROI panel- the colour corresponds to the scalp ROI. The dashed blue line indicates the critical R^2^ threshold based on the distribution

The left fronto-lateral ROI passed the significance threshold from 745 ms until 782 ms (R^2^ range = 0.20 to 0.22) (Figure 7; Figure S15), a period that corresponds to an early CNV component. Within this ROI, a significant curvilinear association between reward-outcome ERP activity and inattentive symptoms was found (β range = 0.36 to 0.56; β threshold (M ±SD) = −.24 ± 0.01) over 48 contiguous samples (741 − 833 ms, *contiguous* p < .001) (Figure 8A; Figure S15). This association was typical-u-shaped. Reward-outcome ERP activity decreased as symptoms transitioned from low to moderate but increased as symptoms transitioned from moderate to high.

**Figure 8.**
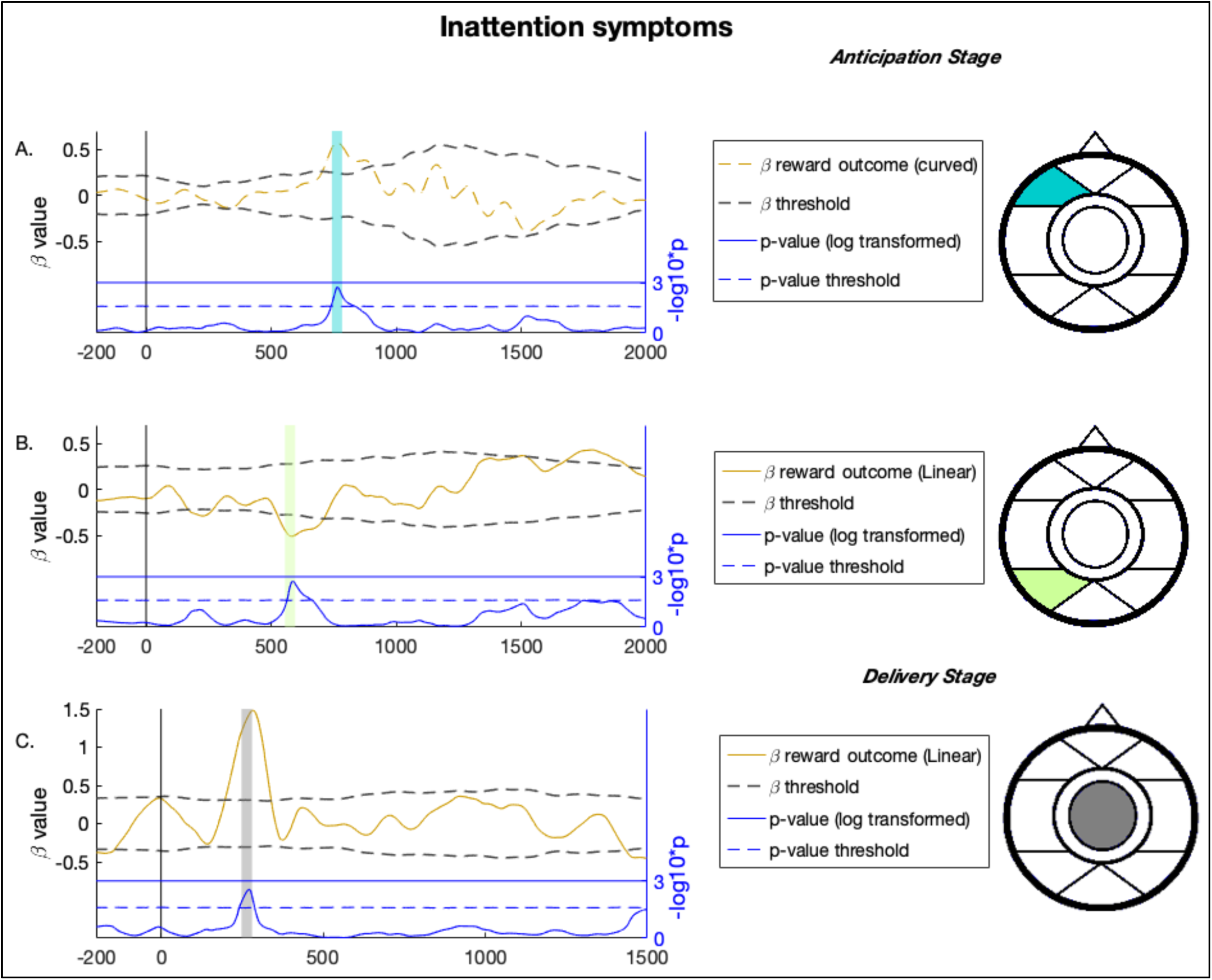
Inattention symptoms were significantly associated with incentive-related ERP activity at 2 scalp ROIs during the anticipation stage and 1 scalp ROI during the delivery stage (left hand side). Significant periods are illustrated by coloured vertical bars in each panel; these coloured bars correspond to the contiguously significant R^2^ values (also in Figure 7 and Figure 9). Further analysis revealed that only certain incentive conditions were associated with inattention symptoms at each scalp ROI. (A) At the fronto-lateral left ROI, the association between ERP activity and symptoms was driven by the reward-outcome (curved association with symptoms). (B) At the posterior-lateral left ROI, the association between ERP activity and symptoms was driven by the reward-outcome (linear association). (C) At the fronto-central ROI, the association between ERP activity and symptoms was driven by the reward-outcome (curved association). Statistical significance was based on the contiguous beta values (coloured dotted lines) that exceeded the bottom 2.5 or top 97.5 percentile of the corresponding beta values from all (7000) null models (i.e. the 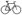threshold; the grey dotted line). Also illustrated are the p-values that corresponds to the beta values. These are presented in blue at the bottom of each panel. We applied the maximum statistic approach to these values to identify p-values that exceeded the top 97.5 percentile of the corresponding p values from all (7000) null models (i.e. a p threshold; the blue dotted line). This value was log transformed so that higher values trended towards statistical significance: where p = .05, −log10(p) = 1.30; where p = .01, −log10(p) = 2; where p = .001, −log10(p) = 3.

The left posterior-lateral ROI passed the significance threshold from 558 ms until 595 ms (R^2^ range = .20 to .22) (Figure 7; Figure S19). Within this ROI, a significant negative association between reward-outcome ERP activity and hyperactive/impulsive symptoms was found (β range = −0.51 to −0.43; β threshold (M ±SD) = −.29 ± 0.01) across 51 contiguous samples (566 − 663 ms, *contiguous* p < .001) (Figure 8B; Figure S19)

#### Delivery Stage

Based on the maximum statistic approach, the critical R^2^ threshold across the threshold was around 0.20 ± 0.005 (M ± SD). The probability of observing two (or more) consecutives samples with a significant R^2^ value was less than one in every 7000 models. Using these criteria, ERP activity from one scalp ROIs was significantly associated with inattentive symptoms (Figure 9). This was the fronto-central ROI.

**Figure 9.**
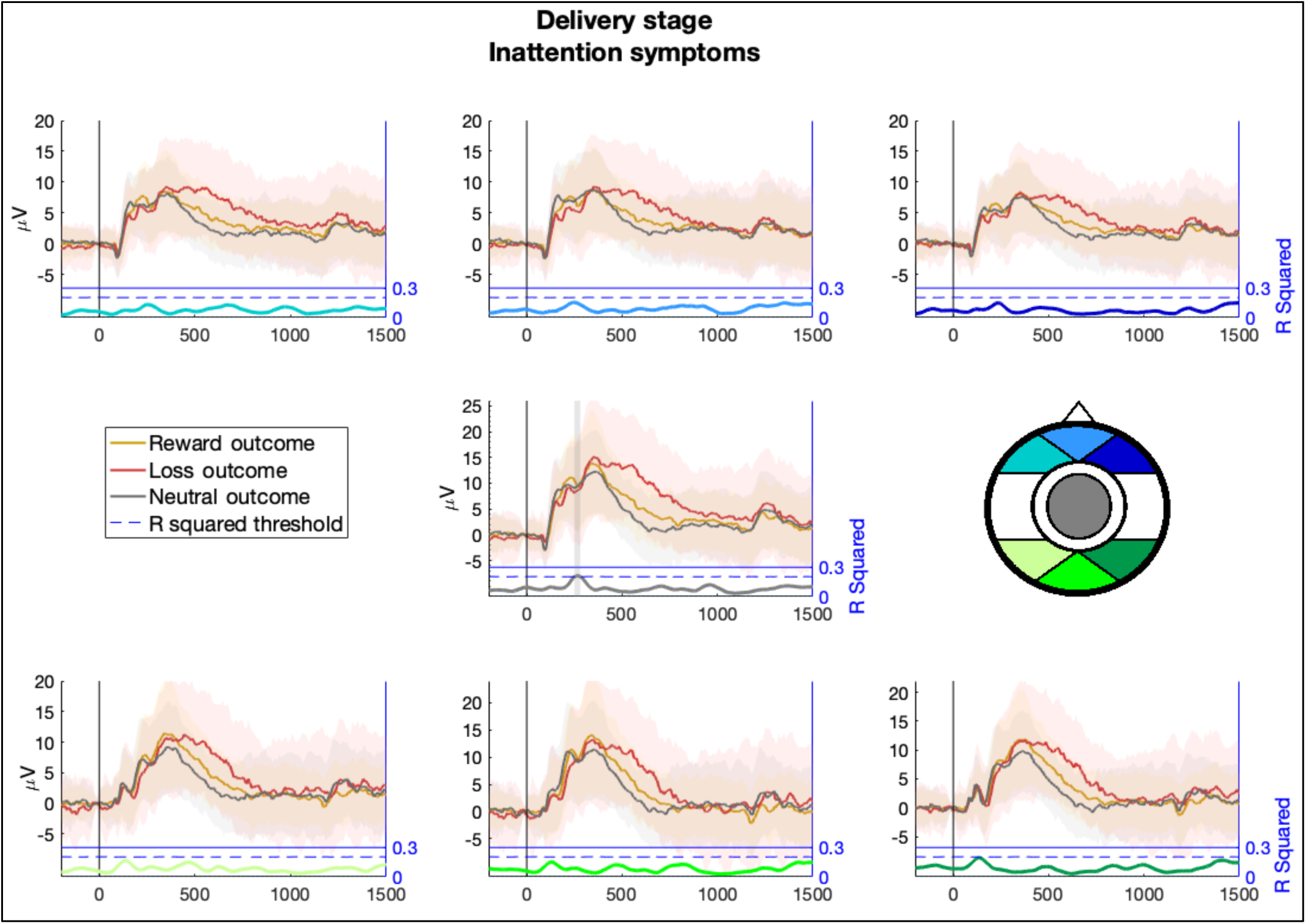
Average ERP amplitudes in response to reward, loss and neutral outcomes during the delivery stage. Each panel represents a scalp ROI. Lighter region surrounding the ERP amplitudes represents the standard deviation. μV = micro volts. The coloured vertical bars illustrate contiguous periods when ERP amplitudes were significantly associated with inattention symptoms. Incentive-related ERP activity at the fronto-central scalp ROI was significantly associated with inattention symptoms. This is based on whether R^2^ values continuously exceeded the top 97.5 percentile of all (7000) null model R^2^ values (i.e. the R^2^ threshold). The R^2^ value at each time point is illustrated at the bottom of each ROI panel- the colour corresponds to the scalp ROI. The dashed blue line indicates the critical R^2^ threshold based on the distribution of null model R^2^ values (i.e. the 97.5 percentile).

The fronto-central ROI passed the significance threshold from 251 ms until 280 ms (R^2^ range = 0.20 to 0.21) (Figure 9; Figure S25). This period corresponds to an ERP component associated with the processing of relative gains; namely, the RewP component. Within this ROI, a significant positive association between reward-related EEG activity and inattentive symptoms was observed (β range = 1.12 to 1.48; β threshold (M ±SD) = .31 ± 0.01) across 25 contiguous samples (243 − 290 ms, *contiguous* p < .001) (Figure 8C; Figure S25).

## Discussion

Reward processing in adult ADHD was explored using the e-MID task. We investigated when, and how, incentive-related ERP activity was associated ADHD symptom dimensions (i.e. hyperactive/impulsive and inattentive symptoms). Specifically, we examined event-related potential (ERP) components that are assumed to reflect brief neuro-cognitive mechanisms involved in the processing, storage and retrieval of reward-relevant information (e.g., cue detection, motor-response preparation and outcome evaluation (19, 21)).

We first investigated associations between incentive-related ERP activity and hyperactive/impulsive symptoms. Cue-P3 amplitudes are linked to early top-down processes including attentional allocation (15, 20–22) as well as the updating of working memory in response to new stimuli (19). Motivationally salient cues exert greater demands on top-down processing and this is reflected by higher cue-P3 amplitude (especially for reward cues) (19, 23). In this study, we found a negative association between cue-P3 amplitudes in response to reward cues (around 200 ms post-cue; left posterior-lateral scalp) and impulsivity symptoms. This association was linear, suggesting that reward anticipation is hypoactive in individuals with elevated hyperactive/impulsive symptoms. This pattern may be characteristic of impaired reward anticipation. If the transfer of brain activity from rewards to their predictive cues is diminished (i.e., impaired reward anticipation), then predictive cues will place fewer demands on top-down processing. As a result, lower cue-P3 potentiation in response to reward-cues would be expected. Importantly, an attentional deficit cannot explain these findings, because cue-P3 amplitudes in response to neutral cues were not significantly associated with symptoms any point in time.

CNV amplitudes in response to reward and loss-cues were significantly associated with hyperactive/impulsive symptoms (around 1500 ms post cue; medial and right posterior-lateral scalp). The relationship between reward-related CNV amplitudes and hyperactive/impulsive symptoms was typical-u-shaped. It is important to note that greater negativity in CNV amplitudes reflects greater motor-response preparation. A typical-u-shaped relation therefore indicates that the extreme ends of hyperactive/impulsive symptoms are associated with smaller CNV amplitudes – that is, poorer motor-response preparation. Interestingly, the opposite pattern was observed for loss-related CNV amplitudes. The association between loss-related CNV amplitudes and hyperactive/impulsive symptoms was an inverted-u-shape. These findings suggest, in the context of motor-response preparation, that the extreme ends of hyperactive/impulsive symptoms are characterised not only by a hyposensitivity to potential rewards but also a hypersensitivity to potential losses. To our knowledge, our study is the first to report this outcome.

We also investigated the relationship between incentive-related ERP activity and inattentive symptoms. CNV amplitudes during reward anticipation were associated with inattentive symptoms. Again, this relationship was u-shaped, suggesting that ADHD symptoms in general are linked to disruptions in incentive-related motor response preparation. Specifically, the extreme ends of ADHD symptomology are associated with hypoactive ERP activity in response to reward-cues during motor response preparation.

Inattention, but not hyperactive/impulsive symptoms, was associated with atypical ERP activity following the delivery of a reward. Specifically, there was a negative association between early ERP amplitudes and inattention (around 250 ms, fronto-central scalp ROI). Recent findings suggest that ERPs at this time and scalp space reflect reward-specific activation elicited from striatal reward regions. This has been referred to as a reward positivity (RewP) component, which positively increases in response to relative gains (24, 25). For example, Becker and colleagues demonstrated higher ERP positive amplitudes for reward relative to loss and neutral outcomes (32). Furthermore, reward-related positivity was specifically associated with a relative increase in VS, mid-cingulate and mid-frontal cortices [34]. While the exact cognitive significance of the RewP component is unclear, its sensitivity to reward outcomes appears to reflect some an aspect of evaluative (or motivational) processing (19). Following on from this, our findings suggest that inattention symptoms, in the context ADHD, are associated with a disruption in the evaluative processing of reward-specific outcomes.

In the current study, ADHD symptom dimensions were associated with specific neuro-cognitive sensitivities during reward processing. Increased hyperactive/impulsive symptoms were associated with hypoactive ERP activity in response to reward-predictive cues; this was evident during initial deployment of attentional resources in the anticipation stage. Increased inattention symptoms were associated with hypoactive ERP activity in response to reward outcomes; this was evident during evaluative processing of outcomes in the delivery stage. Both symptom dimensions were associated with atypical motor-response preparation in response to incentive cues. These findings cannot be explained by a general attentional deficit during task engagement because ERP activity in response to neutral cues was not associated with ADHD symptom dimensions.

Our findings differ from the only other investigation of reward processing in ADHD using the e-MID task. Chronaki and colleagues (15) analysed data from 20 adolescents with ADHD and 26 comparisons, and observed higher cue-P3 amplitude for reward cues in ADHD relative to comparisons. Their finding suggests that reward anticipation is hypersensitive in ADHD. Although the precise reason for these contradicting results is unclear, there are some methodological differences between our and their study. For example, the groups reported in our study are larger and were matched in terms of age, sex, education and IQ. Another key difference between the two studies was the data analysis strategy. Chronaki and colleagues examined between-groups differences in ERP amplitudes at specific ERP time windows. These times were validated through previous research (20). Our analysis was more ‘data-driven’, whereby significant associations between ERP activity and symptoms were extracted across contiguous sample point. Our approach affords a considerable degree of temporal resolution, which may have allowed us to better disentangle the component processes of reward processing and observe novel associations with symptoms. A final difference between the studies was that we examined the association between ERP activity and ADHD symptoms continuously, across the entire sample, rather than between groups (also see (33)). The current approach is therefore commensurate with the emerging view that psychiatric symptoms are dimensional, rather than categorical constructs, with functional impairment occurring at the extreme ends (34, 35).

Some etiological models suggest that ADHD symptoms result from diminished brain activity during reward anticipation (6, 36, 37). If not effectively anticipated, future rewards are less likely to influence immediate actions and decisions. This insensitivity to rewards may manifest as impulsive behavioural traits, including a bias towards larger and more immediate rewards, as well as poorer planning and self-regulation. These models therefore predict irregular brain activity in reward networks during the anticipation stage. Most studies rely on MID tasks with fMRI and, in healthy individuals, increased VS activation is observed during reward relative to neutral cue anticipation (14, 38–40). Adolescent and adult ADHD, in comparison, has been characterized by VS hypoactivity in response to reward-predictive cues (10, 11). These studies also report a negative association between VS activation and impulsive symptoms in ADHD. However, others have found no evidence of abnormal brain activity during reward anticipation in ADHD. For example, Stoy and colleagues observed significant VS activation during reward anticipation but no differences between adult ADHD and comparisons (17). Paloyelis and colleagues also demonstrated that VS activation during reward anticipation did not differ between an adult ADHD group and comparisons (16). Also, von Rhein and colleagues observed an increase in VS activation during reward anticipation but no differences between a large group of adolescents and adults with ADHD versus comparisons (18).

Although our analysis was ERP-based, the results are relevant to the previous functional neuroanatomy literature. Pfabigan and colleagues administered a MID task to healthy individuals twice, first with fMRI and again with EEG (22). They reported a positive association between reward-related cue-P3 amplitudes and reward-related VS activation, but no association between CNV amplitudes and VS activation. In this study, posterior cue-P3 and CNV amplitudes in response to rewards were atypical in cases of elevated impulsivity. Therefore, the negative association between reward-related cue-P3 amplitudes and impulsivity may reflect VS hypoactivity during reward anticipation in ADHD. As such, our study supports previous evidence of lower VS activity during reward-anticipation in ADHD (9-11, 41, 42). The current findings also contribute to the literature by demonstrating an association between inattentive symptoms and hypo-sensitivity to rewards during evaluative processing of outcomes.

A limitation of the current study is that we cannot speak to the causal connections between different neuro-cognitive mechanisms. One possibility is that a weaker allocation of attentional resources to reward cues during the anticipation stage (a correlate of impulsivity) is a consequence of the inefficient evaluation of reward outcomes during the delivery stage (a correlate of inattention). However, the reverse could also be true. For example, an ‘aversion-of-delay’ model of ADHD posits that deficits in the ability to anticipate rewards will accrue inattentive symptoms across development (36, 37). To parse these causal connections, future research will benefit from larger group sizes with a community sample that includes participants who are characterised by just one of ADHD symptom dimensions (e.g. distinct hyperactive/impulsive and inattentive subgroups).

In conclusion, hyperactive/impulsive and inattentive symptoms are associated with hyposensitivity to rewards during the anticipation and delivery stage. Increased hyperactive/impulsive symptoms were associated with hypoactive ERP activity in response to reward-predictive cues; this was evident around the time of a cue-P3 component. Increased inattention symptoms were associated with hypoactive ERP activity in response to reward outcomes; this was evident around the time of a RewP component. Finally, when anticipating rewards and losses, CNV amplitudes were reduced at the extreme ends of hyperactive/impulsive symptoms and inattention symptoms. These results shed new light on the relationship between the neurological mechanisms of reward processing and symptom dimensions underlying ADHD.

## Acknowledgements

We would like to thank Ken Kilbride and ADHD Ireland for their valuable support throughout this project. This work is supported by the Irish Research Council Government of Ireland Postdoctoral Fellowship grants-awarded to M. Bennett (GOIPD/2016/617) and H. Kiiski (grant number GOIPD/2015/777). M. Bennett is also partially supported by the Wellcome Trust (104908/Z/14/Z). Z. Chao is supported by China Scholarship Council. F. Farina is supported by an Irish Research Council Enterprise Partnership Postdoctoral Award (EPSPD/2017/110). D. Roddy is supported by Irish Health Research Board grant to the REDEEM group at Trinity College Institute of Neuroscience and Department of Psychiatry (201651.12553). C. Kelly was supported by the Trinity College Dublin Pathfinder seed grant. R. Whelan is supported by Brain & Behavior Research Foundation’s NARSAD Young Investigator grant (23599) and a Science Foundation Ireland grant (16/ERCD/3797). The other authors did not have funding to disclose. The authors had no conflicts of interest. The study sponsors had no involvement in the collection, analysis and interpretation of data or in the writing of the manuscript.

## Notes

https://osf.io/k4hqz/

