## Supplementary Materials for "Hyperactive/impulsive and inattention symptoms are associated with reduced ERP activity during different reward processing stages: Evidence from the electrophysiological Monetary Incentive Delay Task in adult ADHD"

**Authors:** M. P. Bennett[1,2]* PhD, H. Kiiski[1] PhD, Z. Cao[3] MSc, F. Farina[1] PhD, R. Knight[1] MSc, A. Sweeney[1] MSc, D. Roddy[1,4] MD, C. Kelly[1,4] PhD & R. Whelan[1,5]* PhD

**Affiliations:**

1 Trinity College Institute of Neuroscience, Trinity College Dublin, Dublin, Ireland.

2 MRC- Cognition and Brain Science Unit, University of Cambridge, UK.

3 School of Psychology, University College Dublin

4 Department of Physiology, School of Medicine, University College Dublin, Dublin 4, Ireland.

5 Global Brain Health Institute, Trinity College Dublin, Dublin, Ireland.

*** To whom correspondence should be addressed**

Robert Whelan-

or

Marc Bennett- mailto:

Trinity College Institute of Neuroscience, Trinity College Dublin, College Green, Dublin 2, Ireland.

**Keywords:** Attention Deficit/Hyperactivity Disorder, Reward processing, e-MID task, reinforcement learning, EEG.

**Data collection Protocol**

Participants first completed a telephone interview to establish inclusion/exclusion criteria. An online questionnaire battery was then completed, which included demographic information, Conners' Adult ADHD Rating Scales (CAARS) (1), state-trait anxiety inventory (STAI) (2) and the NEO personality inventory (3). This was administered using Qualtrics (https://www.qualtrics.com/) and SurveyCTO (https://www.surveycto.com). The questionnaire battery was completed approximately one week before or after EEG testing. On the day of testing, participants completed a Structured Clinical Interview of the DSM-IV to screen for additional exclusion criteria (Table S1). IQ was assessed using the National Adult Reading Test (NART) (4) and then age-corrected (5, 6). Afterwards, EEG testing began as described in the main text.

**Additional self-report measurements**

**The Neuroticism-Extraversion-Openness Five Factor Inventory** (NEO-FI) (3) assessed personality traits, including individual differences on five subscales for Neuroticism, Extraversion, Openness, Agreeableness and Conscientiousness. Each subscale comprised 12 items on a 5-point Likert scale (0=strongly disagree; 4=strongly agree). Internal consistency for the five domains is good (α = .68 to .86;) (3).

**State Trait Anxiety Inventory** (2). The STAI is a 40 item self-report measure of trait and state anxiety; higher scores indicate higher levels of anxiety (7). The

STAI has good test-retest reliability (coefficients range from 0.31 to 0.86) and strong internal consistency (α = 0.86 for young adults).

**Conner's Adult ADHD Rating Scale** (1). The CAARS comprises four sub-scales of ADHD symptomatology: inattention, hyperactivity/impulsivity and self-concept. Psychometric analysis of self-reported CAARS reveals strong internal reliability (average α = .75) and test–retest correlations (r = 0.80) (8). Also, it is a valid measure that correlates moderately well with other measures of ADHD (e.g. Wender Utah Ratings Scale; *r* inattention = 0.37, *r* hyperactivity = 0.48, *r* impulsitvity = 0.67 and self-concept = 0.37).

**EMID Tracking algorithm**

The tracking-algorithm successfully established a response hit rate of ~66% in all conditions (see Table S2).

### Additional demographic details

*Table S1. Additional demographic details and Structural Clinical Interview for the DSM*

| *Demographic Items* | Total (N = 68) | | ADHD (N = 32) | | Comparison (N = 36) | | | | |
|---|---|---|---|---|---|---|---|---|---|
| | % | | % | | % | | $x2$ | df | p |
| **Monthly Income** | | | | | | | 3.78 | 3 | .29 |
| ≤ €900 | 73.50 | | 78.10 | | 69.40 | | | | |
| €900-€1350 | 8.80 | | 3.10 | | 13.90 | | | | |
| €1350-€1800 | 8.80 | | 6.30 | | 11.10 | | | | |
| ≥ €1800 | 8.80 | | 12.50 | | 5.60 | | | | |
| **Profession** | | | | | | | 8.65 | 6 | 0.19 |
| Student | 58.80 | | 56.30 | | 61.10 | | | | |
| Unemployed | 10.30 | | 18.80 | | 2.80 | | | | |
| Unskilled | 1.50 | | 0.00 | | 2.80 | | | | |
| Semi-Skilled | 5.90 | | 3.10 | | 8.30 | | | | |
| Skilled, clerical/sales | 4.40 | | 0.00 | | 8.30 | | | | |
| Semi-professional, managers, technician | 5.90 | | 6.30 | | 5.60 | | | | |
| Professional | 11.80 | | 12.50 | | 11.10 | | | | |
| | **M** | **SD** | **M** | **SD** | **M** | **SD** | **T** | **df** | **p** |
| **Age of ADHD Diagnosis** | | | 19.96 | 10.94 | | | | | |
| **Conners ADHD sub-scales** | | | | | | | | | |
| ADHD Risk Score | 57.96 | 12.67 | 65.81 | 9.71 | 50.97 | 10.83 | 5.92 | 66 | <.001 |
| Hyperactivity/Restlessness | 54.06 | 10.95 | 59.28 | 9.11 | 49.42 | 10.45 | 4.13 | 66 | <.001 |
| Inattention/Memory | 60.53 | 15.14 | 71.06 | 12.19 | 51.17 | 10.75 | 7.15 | 66 | <.001 |
| Impulsivity/Emotionality | 54.62 | 13.56 | 60.94 | 11.17 | 49 | 13.14 | 4.01 | 66 | <.001 |
| Self-Concept | 54.68 | 10.85 | 58.34 | 10.91 | 51.42 | 9.82 | 2.76 | 66 | .008 |
| **Structured Clinical Interview (SCID[1])** | **%** | **(N)** | **%** | **(N)** | **%** | **(N)** | **x2** | **df** | **p** |
| ***Mood Disorder*** | | | | | | | | | |
| Mood Disorders | 7.94 | (5/63) | 16.67 | (5/30) | 0.00 | (0/33) | 5.97 | 1 | .02* |
| Dysthymia | 9.52 | (6/63) | 20.00 | (6/30) | 0.00 | (0/33) | 7.3 | 1 | .007 |
| ***Eating Disorder*** | | | | | | | | | |
| Anorexia Nervosa | 3.17 | (2/63) | 3.33 | (1/30) | 3.03 | (1/33) | 0.005 | 1 | 0.95 |
| Bulimia Nervosa | 11.11 | (7/63) | 21.21 | (7/33) | 0.00 | (0/33) | 8.66 | 1 | .003 |
| ***Substance Abuse*** | | | | | | | | | |
| Alcohol Abuse | 12.69 | (8/63) | 20.20 | (6/30) | 6.06 | (2/33) | 2.75 | 1 | .10 |
| Non-alcohol Abuse | 26.98 | (17/63) | 50.00 | (15/30) | 6.06 | (2/33) | 15.39 | 1 | <.001 |
| ***Anxiety Disorder Screening Items*** | | | | | | | | | |
| Panic Disorder | 7.90 | (5/63) | 43.33 | (13/30) | 6.06 | (2/33) | 12.03 | 1 | <.001 |
| Generalized Anxiety Disorder | 17.50 | (11/63) | 30.00 | (9/30) | 6.06 | (2/33) | 6.25 | 1 | .01 |
| Social Anxiety Disorder | 27.00 | (17/63) | 36.67 | (11/30) | 18.18 | (6/33) | 2.72 | 1 | .10 |
| Specific Phobia | 1.60 | (1/63) | 23.33 | (7/30) | 0.00 | (0/33) | 9.35 | 1 | .009 |
| OCD (thoughts) | 9.50 | (6/63) | 13.33 | (4/30) | 6.06 | (2/33) | .97 | 1 | .33 |
| OCD (compulsions) | 9.50 | (6/63) | 16.67 | (5/30) | 3.03 | (1/33) | 3.39 | 1 | .07 |
| PTSD (experienced trauma) | 23.70 | (15/63) | 26.67 | (8/30) | 9.09 | (3/33) | 3.37 | 1 | .07 |

1: An abreviated SCID was introduced later in data collection (i.e. 30 ADHD and 33 controls). Also, a later version of the SCID included screening items for anxiety disorders (i.e. 12 ADHD and 15 controls). * p ≤ .05. ** p ≤ .01 *** p ≤ .001.

*Table S2 Target hit-rate per condition*

| | Total | | ADHD | | Comparison | | | | |
|---|---|---|---|---|---|---|---|---|---|
| *Condition* | **M** | **SD** | **M** | **SD** | **M** | **SD** | **T** | **df** | **p** |
| **Reward cue (%)** | 68.57 | 7.57 | 68.44 | 6.62 | 68.68 | 8.42 | 0.13 | 66 | .90 |
| **Loss cue (%)** | 69.11 | 9.37 | 70.16 | 6.93 | 68.18 | 11.11 | 0.87 | 66 | .39 |
| **Neutral cue (%)** | 66.12 | 7.17 | 66.09 | 6.47 | 66.15 | 7.85 | 0.03 | 66 | .97 |

**EEG Pre-processing and data loss**

EEG data were bandpass filtered between 0.1 and 95 Hz, with a notch filter at 48-52 Hz. Data were then separated into epochs of 200 ms before and 2000 ms after cue/feedback onset. FASTER identified and extracted artefactual (non-neural) independent components, removed epochs containing large artefacts (muscle twitches and eye-blinks) and interpolated channels with poor signal quality. Visual inspection of the data was then performed using ManualQC plug-in (https://github.com/zh1peng/EEGQC_GUI) and additional epochs were removed where necessary. A mean (±SD) of 15.26 (±12.78) % of trials were removed during pre-processing. There was no differences in data loss between those with an ADHD diagnosis (13.82 (±10.12)%) and comparisons (16.53 (±14.78)%), $t(df)$ = .87(66), $p$ =.39. Preprocessing scripts are available online (https://osf.io/k4hqz/)..

**Regression models at each scalp ROI.**

At each scalp ROI, regression models were calculated (one at each individual time point) to determine when ERP activity was associated with hyperactive/impulsive and inattention symptoms. Scalp ROIs were selected for further inspection when the R squared value was deemed significant. This was based on a maximum statistic approach- R squared values were required to continuously exceed the top 97.5 percentile of 7000 null model (one from each scalp ROI) $R^2$ values (i.e. $R^2$ threshold). At this point, the contribution of individual incentive-related ERP predictors was examined. This was accomplished by examining the beta values for reward, loss and neutral terms. Again, statistical significance was based on the contiguous beta values that exceeded the bottom 2.5 or top 97.5 percentile of the corresponding beta values from the 7000 null models (i.e. the $\beta$ threshold). Although only the main effects are described in the manuscript, the findings for each regression model at each ROI are illustrated below (Figures S1-S28). These figures can be described as follows:

In each figure, $R^2$ values are illustrated in the panel in the bottom left-hand corner (purple line) as is the critical $R^2$ threshold (dark solid line). The position of the scalp space is illustrated in the panel in the bottom right-hand corner. The other left-hand side panels illustrate the beta values for incentive-related ERP activity (reward = gold, loss = red, and neutral = blue). The solid coloured lines represent terms that estimate a linear association between ERP activity and symptoms (i.e. EPR). The dashed coloured lines represent terms that estimate a curvilinear association between ERP activity and symptoms (i.e. $ERP^2$). In these panels, the dark lines represent the critical beta threshold (solid = linear terms; dashed = curved terms). The remaining, right-hand side illustrate the p-values that correspond to these beta values. (Again, reward = gold, loss = red, and neutral = blue). The solid coloured lines represent a linear association between ERP activity and symptoms. The dashed coloured lines represent a curved association between ERP activity and symptoms. We also applied the maximum statistic approach to these values to identify p-values that exceeded the top 97.5 percentile of the corresponding p values from all null models. Here, the dark lines represent the critical beta threshold (solid = linear terms; dashed = curved terms). This value was log transformed so that higher values trended towards statistical significance: where $p = .05$, $-\log10(p) = 1.30$; where $p = .01$, $-\log10(p) = 2$; where $p = .001$, $-\log10(p) = 3$.

Data analysis scripts are available online (https://osf.io/k4hqz/)..

**Hyperactive/impulsive symptoms and anticipation stage ERP activity at fronto-lateral left ROI**

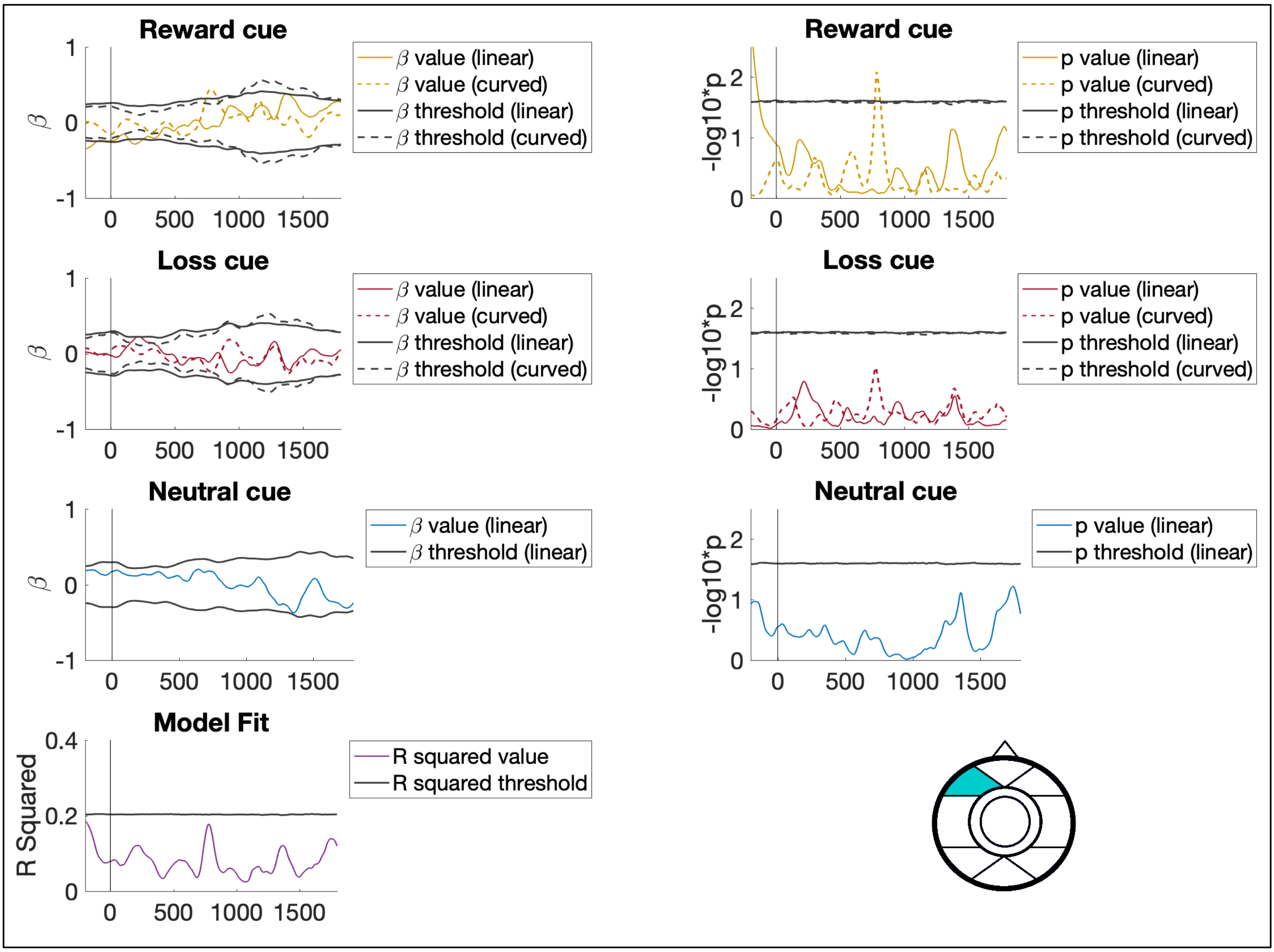


Figure S1. Association between anticipation stage ERP activity and hyperactive/impulsive symptoms at the fronto-lateral left scalp ROI.

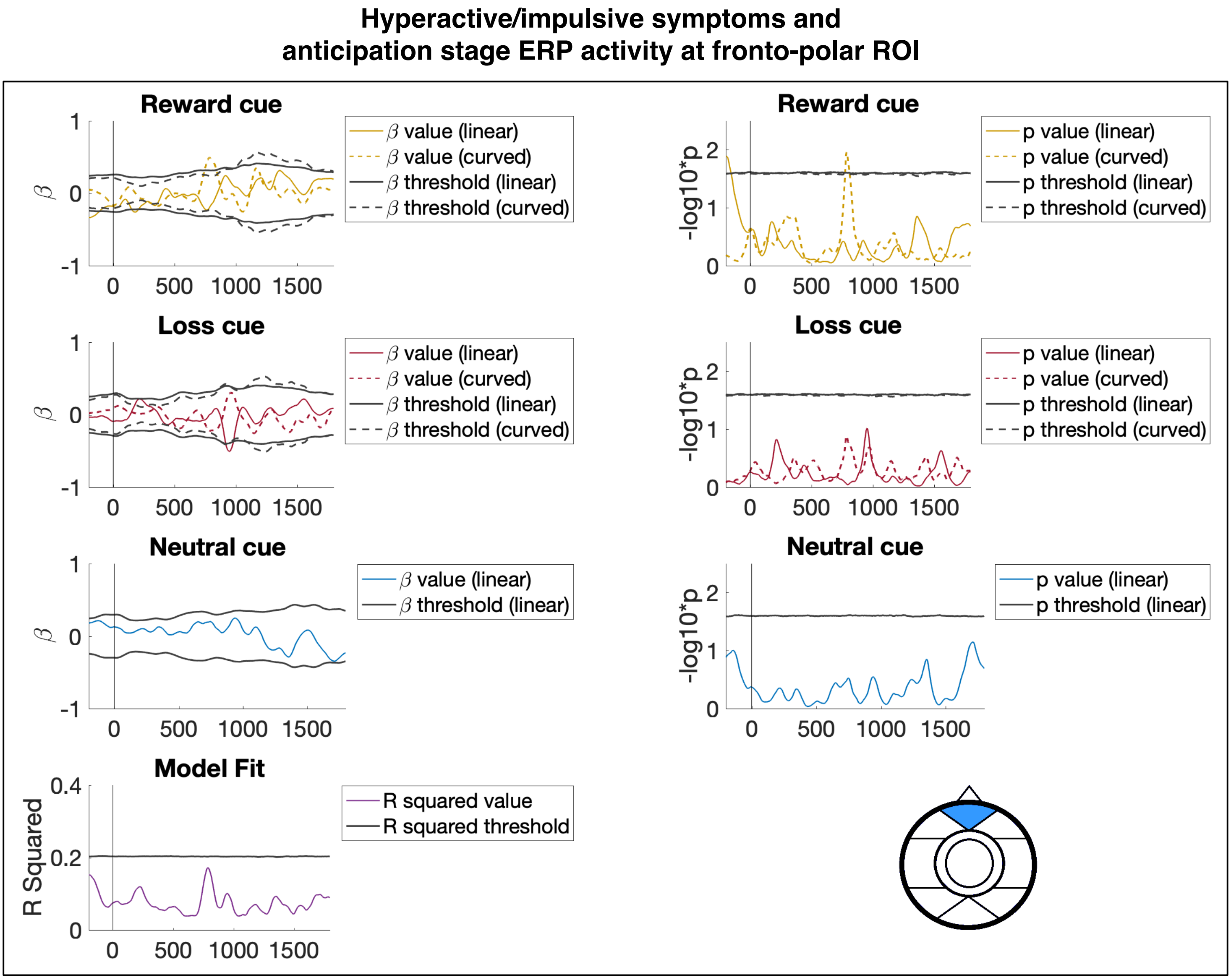


Figure S2. Association between anticipation stage ERP activity and hyperactive/impulsive symptoms at the fronto-polar scalp ROI.

**Hyperactive/impulsive symptoms and anticipation stage ERP activity at fronto-lateral right ROI**

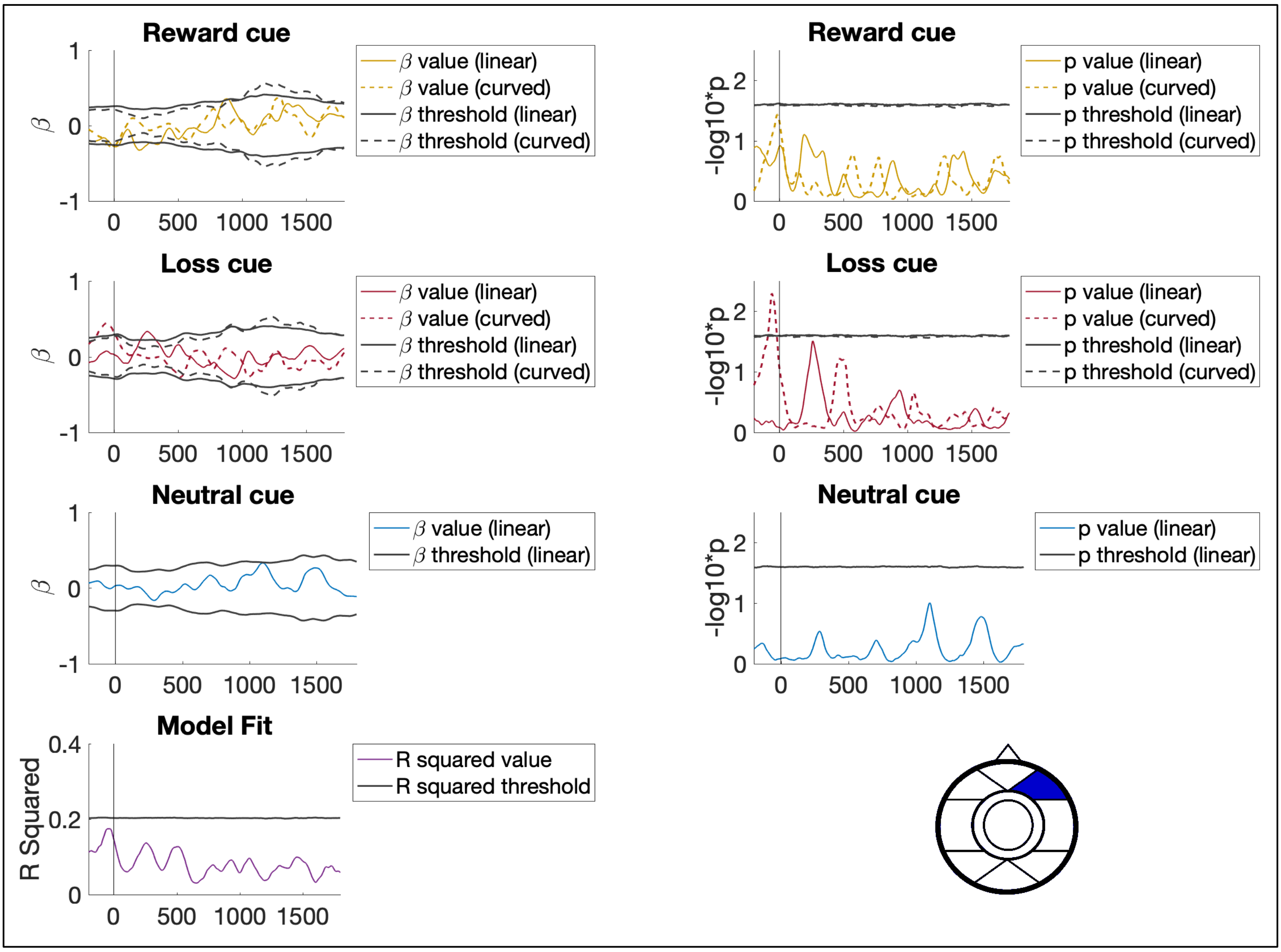


Figure S3. Association between anticipation stage ERP activity and hyperactive/impulsive symptoms at the fronto-lateral right scalp ROI.

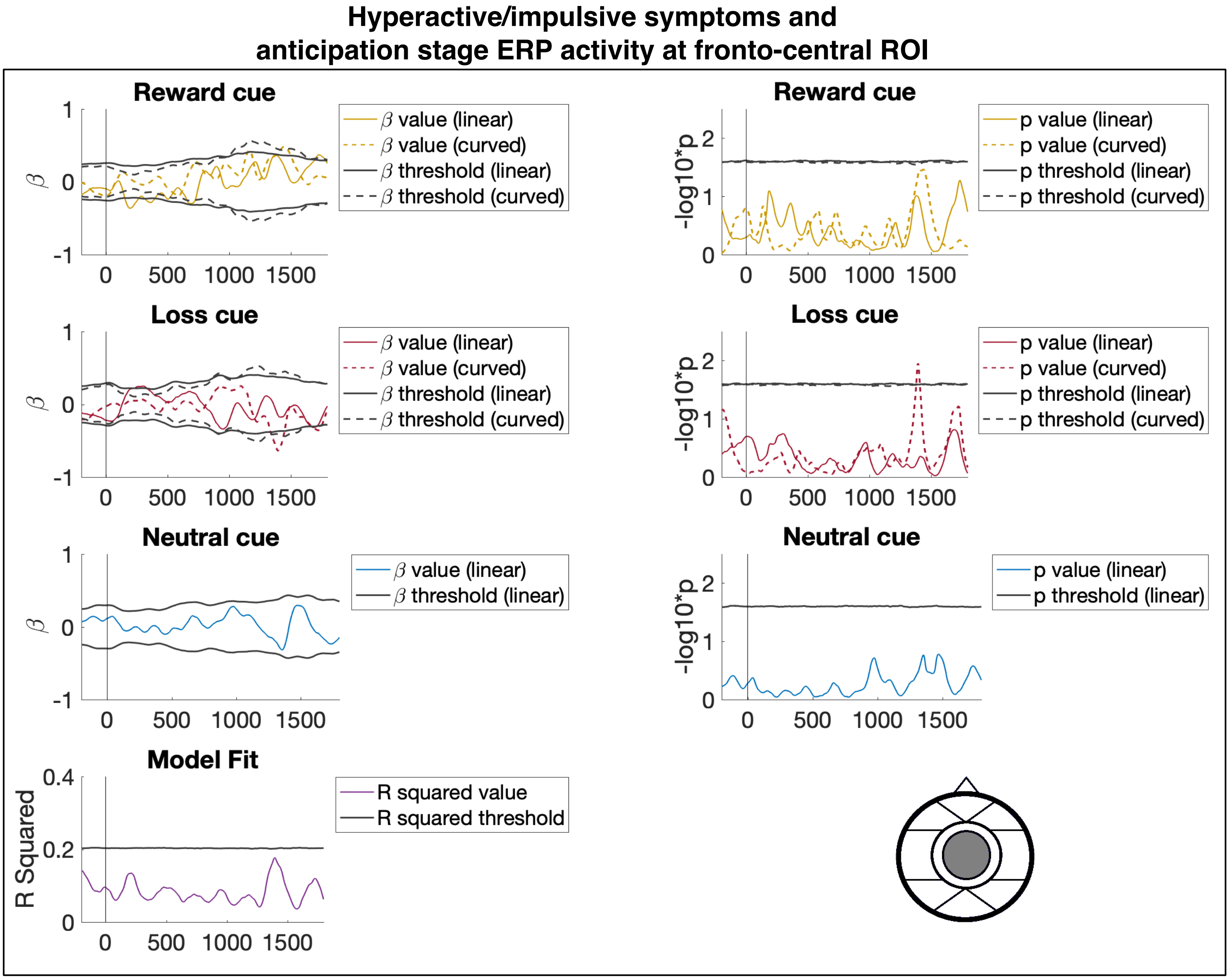


Figure S4. Association between anticipation stage ERP activity and hyperactive/impulsive symptoms at the fronto-central scalp ROI.

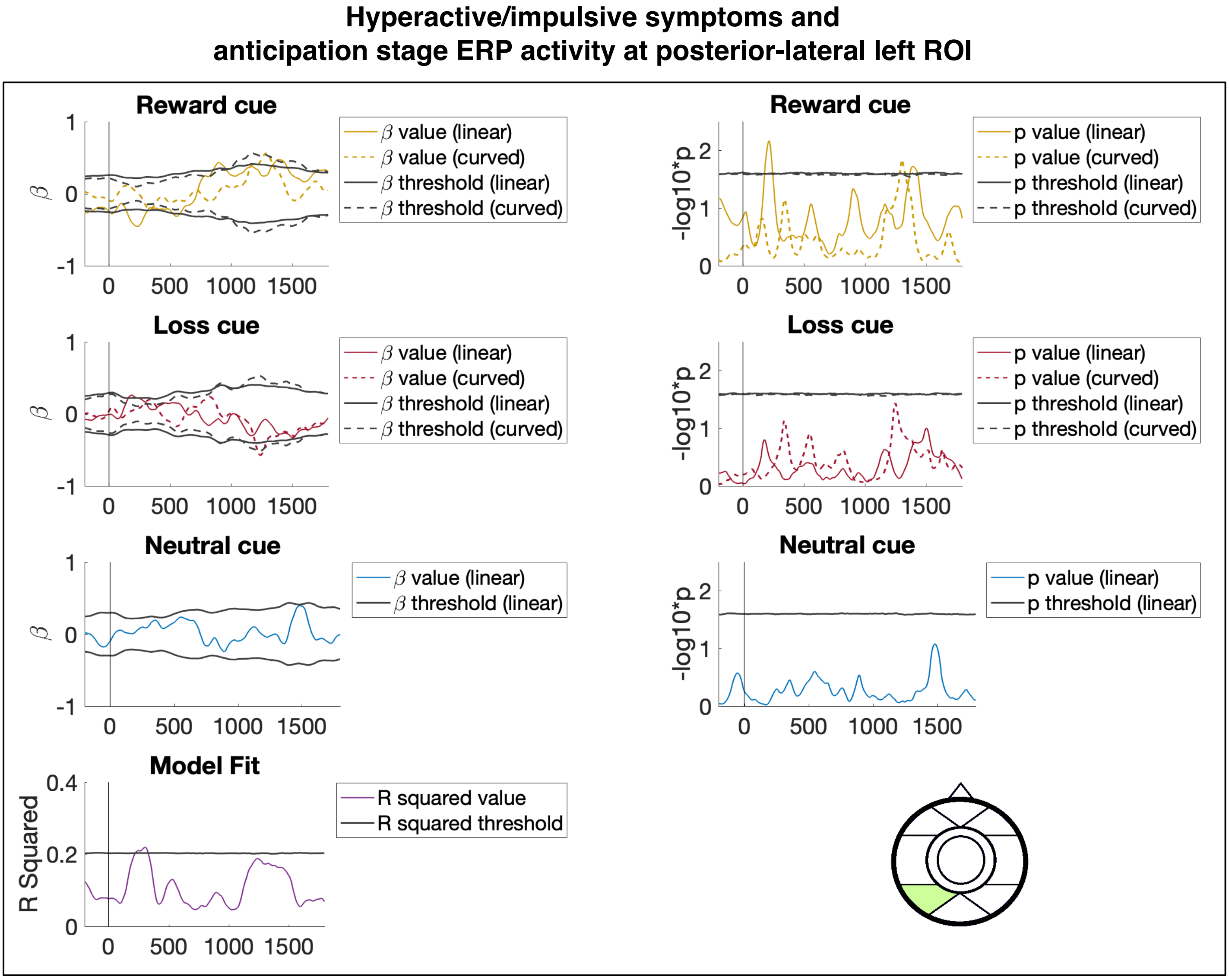


Figure S5. Association between anticipation stage ERP activity and hyperactive/impulsive symptoms at the posterior-lateral left scalp ROI.

**Hyperactive/impulsive symptoms and**
**Anticipation stage ERP activity at occipital-medial ROI**

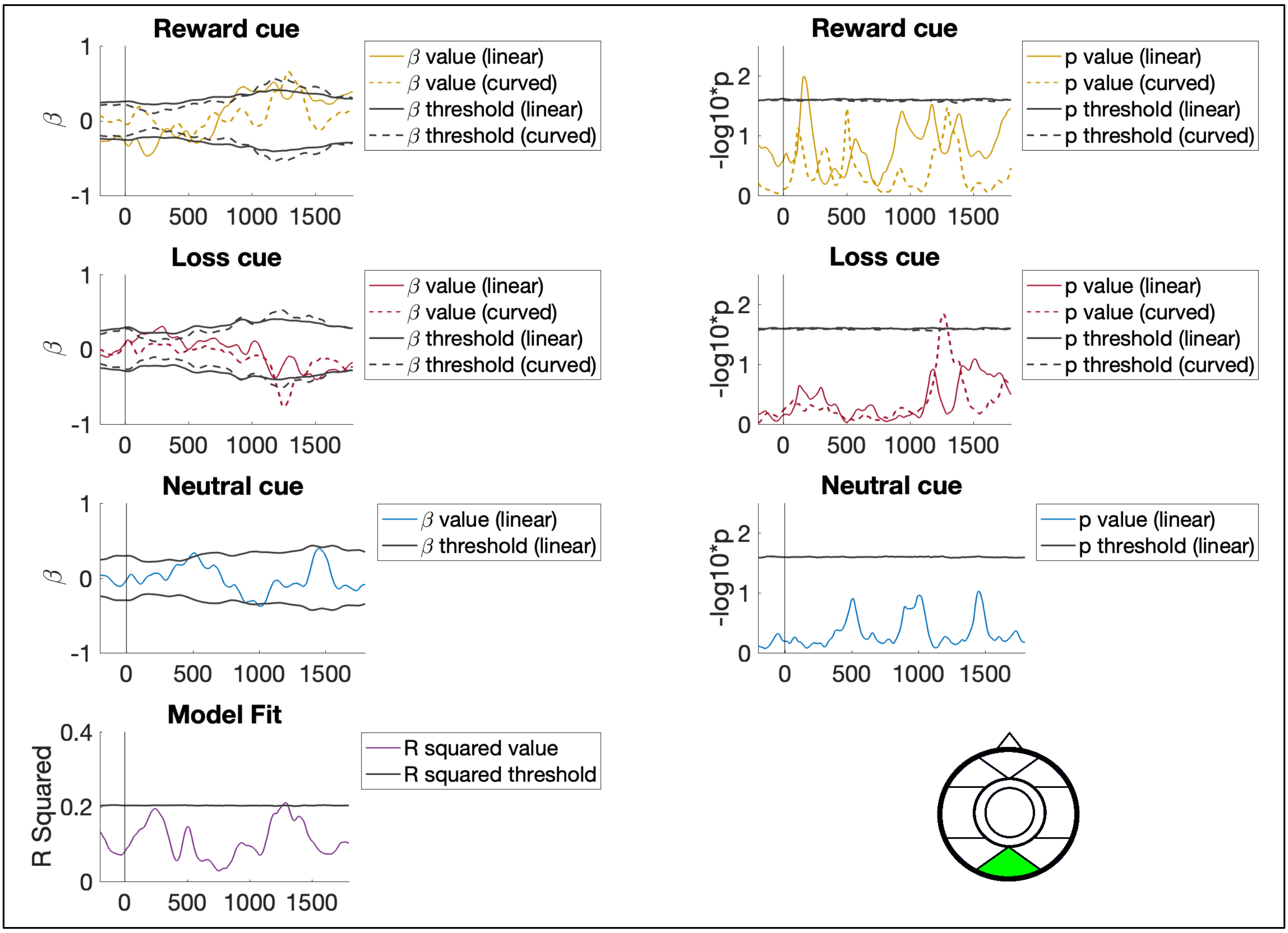


Figure S6. Association between anticipation stage ERP activity and hyperactive/impulsive symptoms at the occipital-medial scalp ROI.

**Hyperactive/impulsive symptoms and anticipation stage ERP activity at posterior-lateral right ROI**

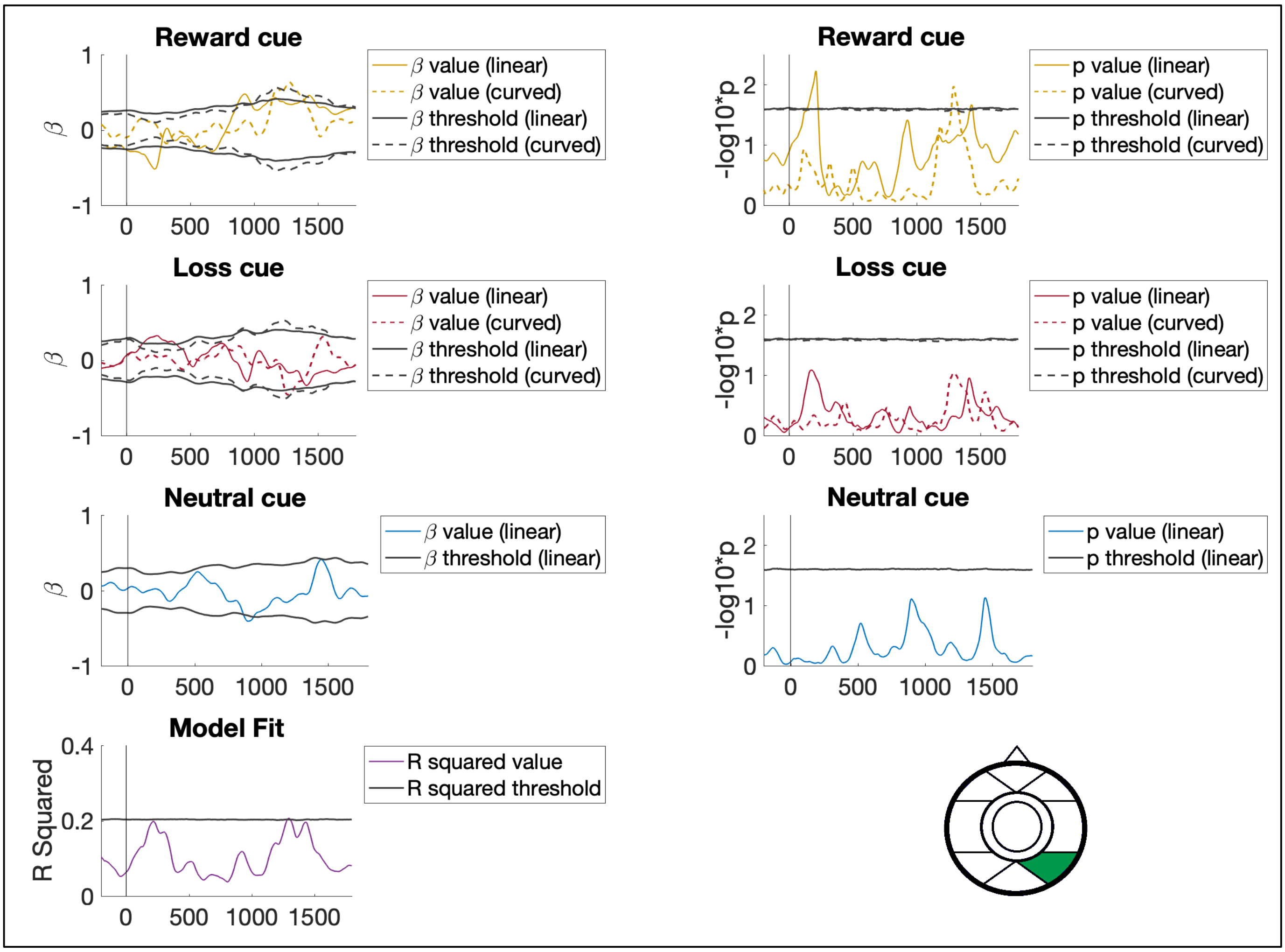


Figure S7. Association between anticipation stage ERP activity and hyperactive/impulsive symptoms at the posterior-lateral right scalp ROI.

**Hyperactive/impulsive symptoms and delivery stage ERP activity at fronto-lateral left scalp ROI**

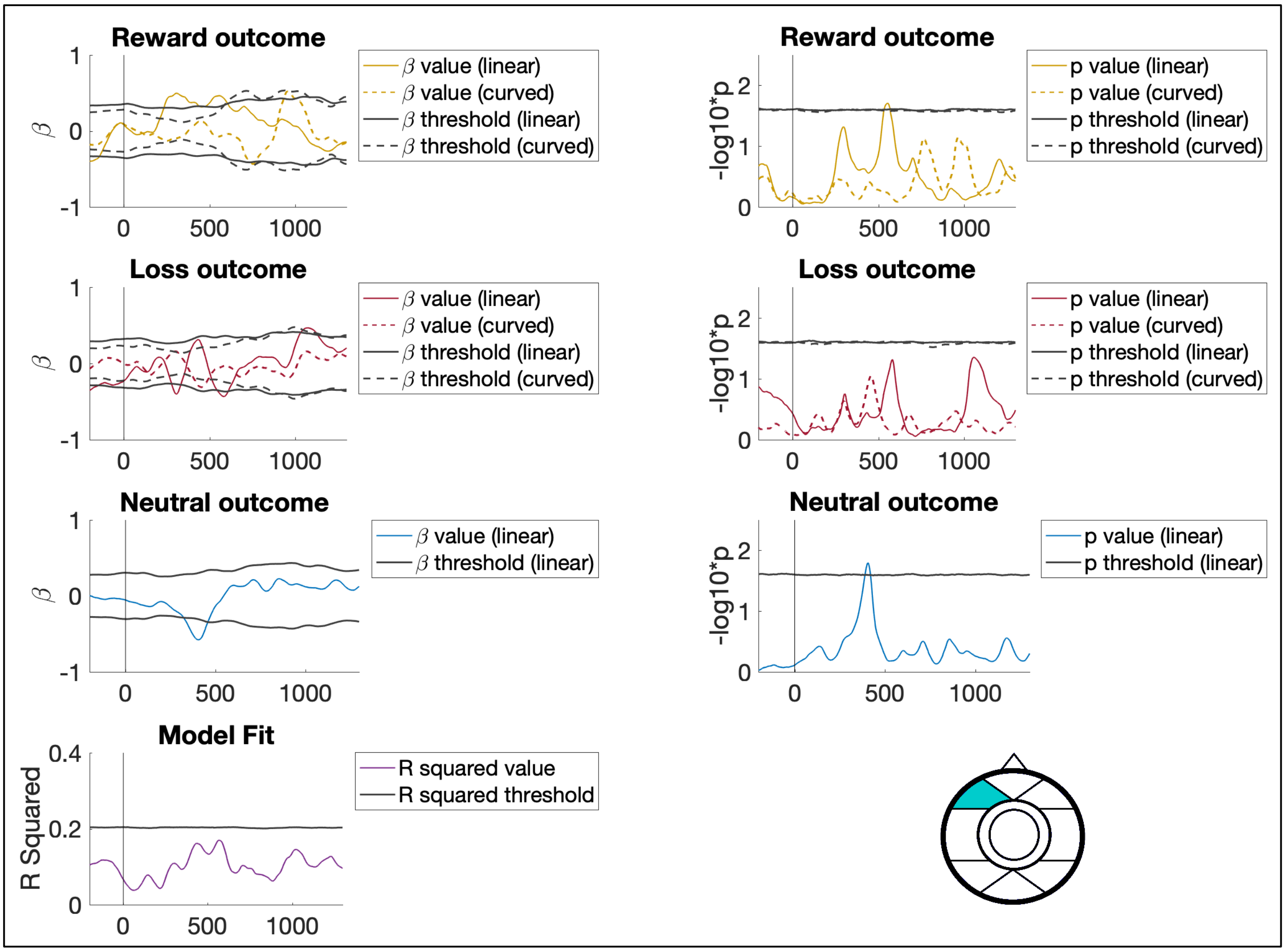


Figure S8. Association between delivery stage ERP activity and hyperactive/impulsive symptoms at the fronto-lateral left scalp ROI.

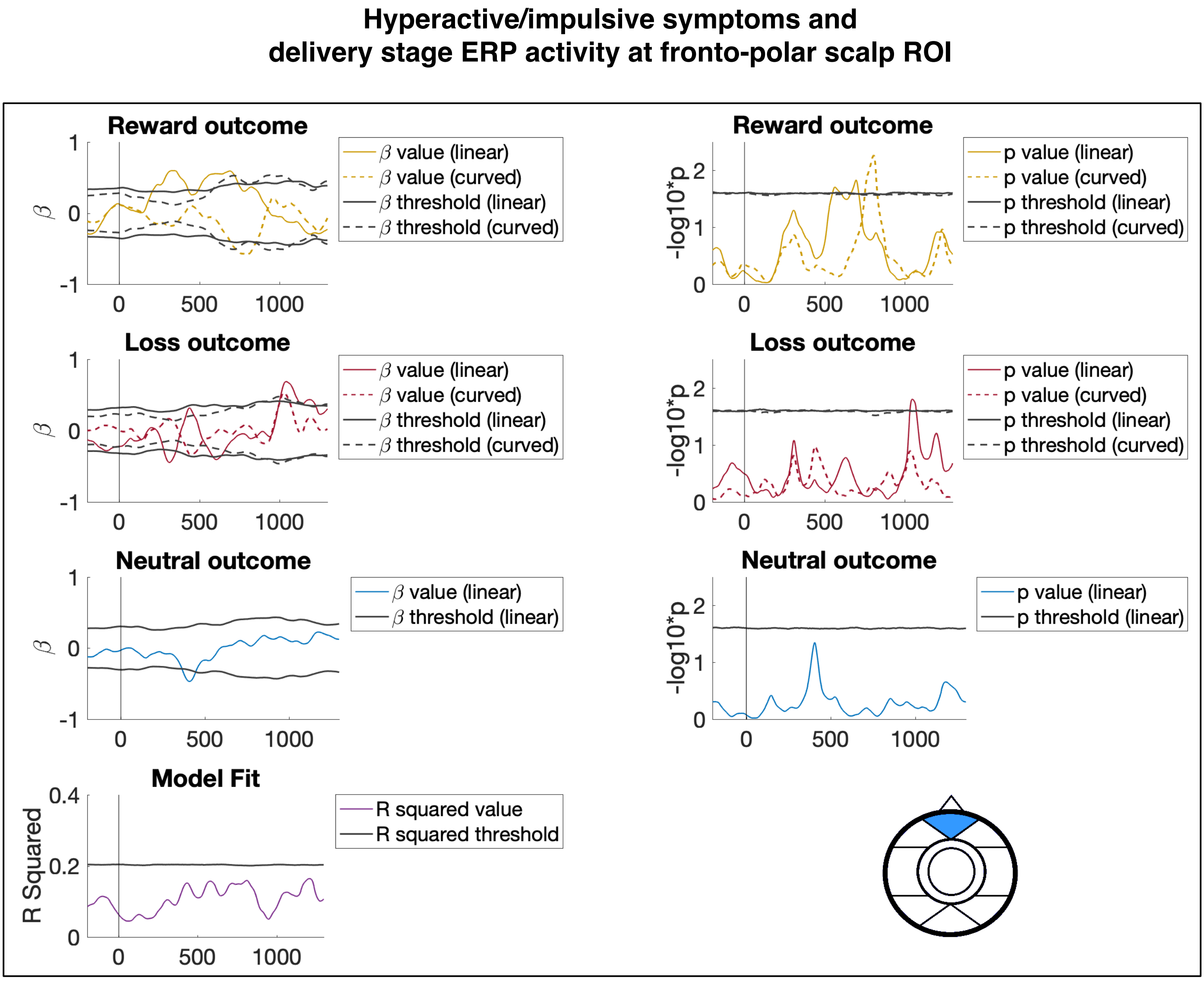


Figure S9. Association between delivery stage ERP activity and hyperactive/impulsive symptoms at the fronto-polar scalp ROI.

**Hyperactive/impulsive symptoms and delivery stage ERP activity at fronto-lateral right scalp ROI**

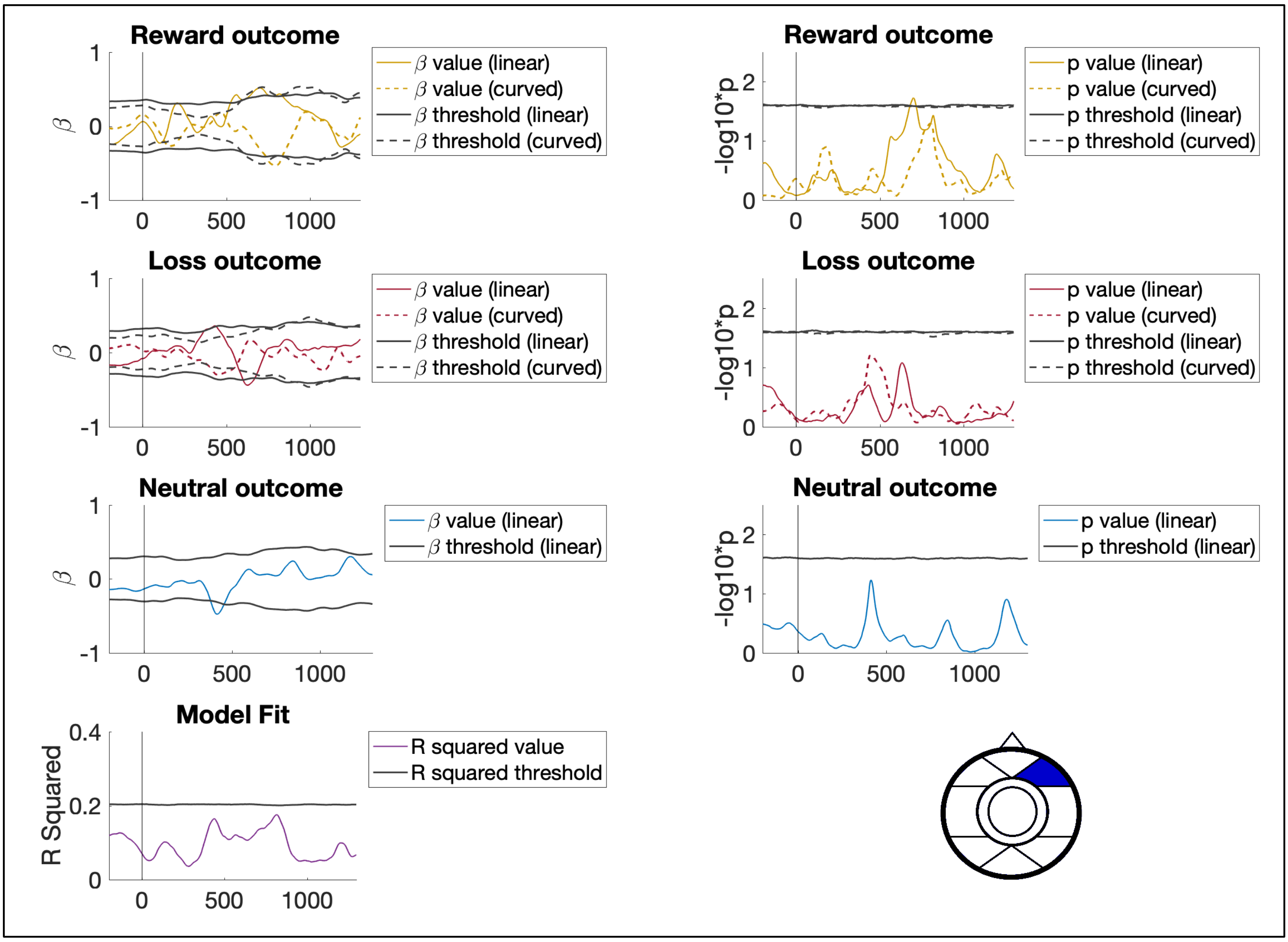


Figure S10. Association between delivery stage ERP activity and hyperactive/impulsive symptoms at the fronto-lateral right scalp ROI.

**Hyperactive/impulsive symptoms and delivery stage ERP activity at fronto-central scalp ROI**

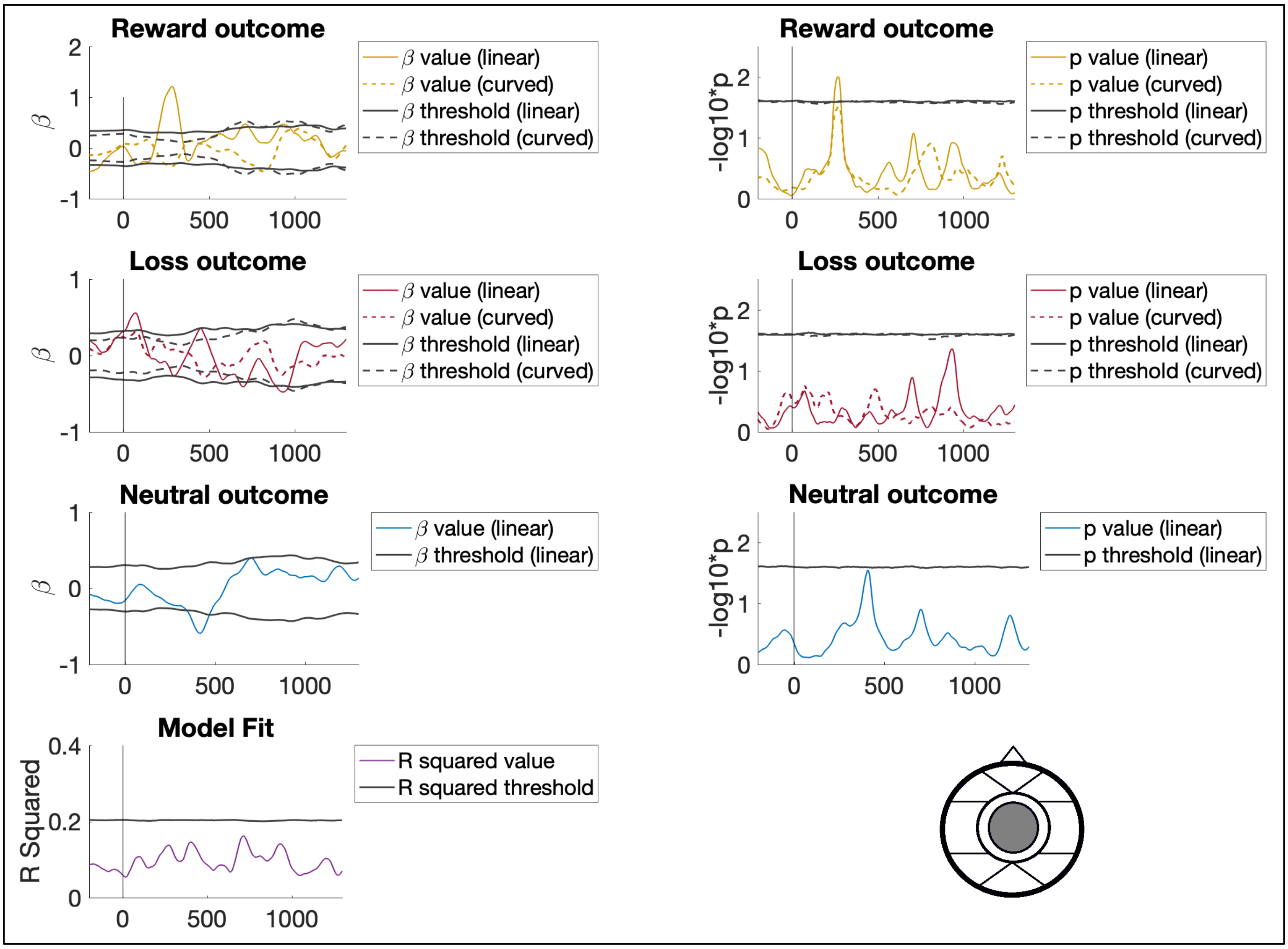


Figure S11. Association between delivery stage ERP activity and hyperactive/impulsive symptoms at the fronto-central scalp ROI.

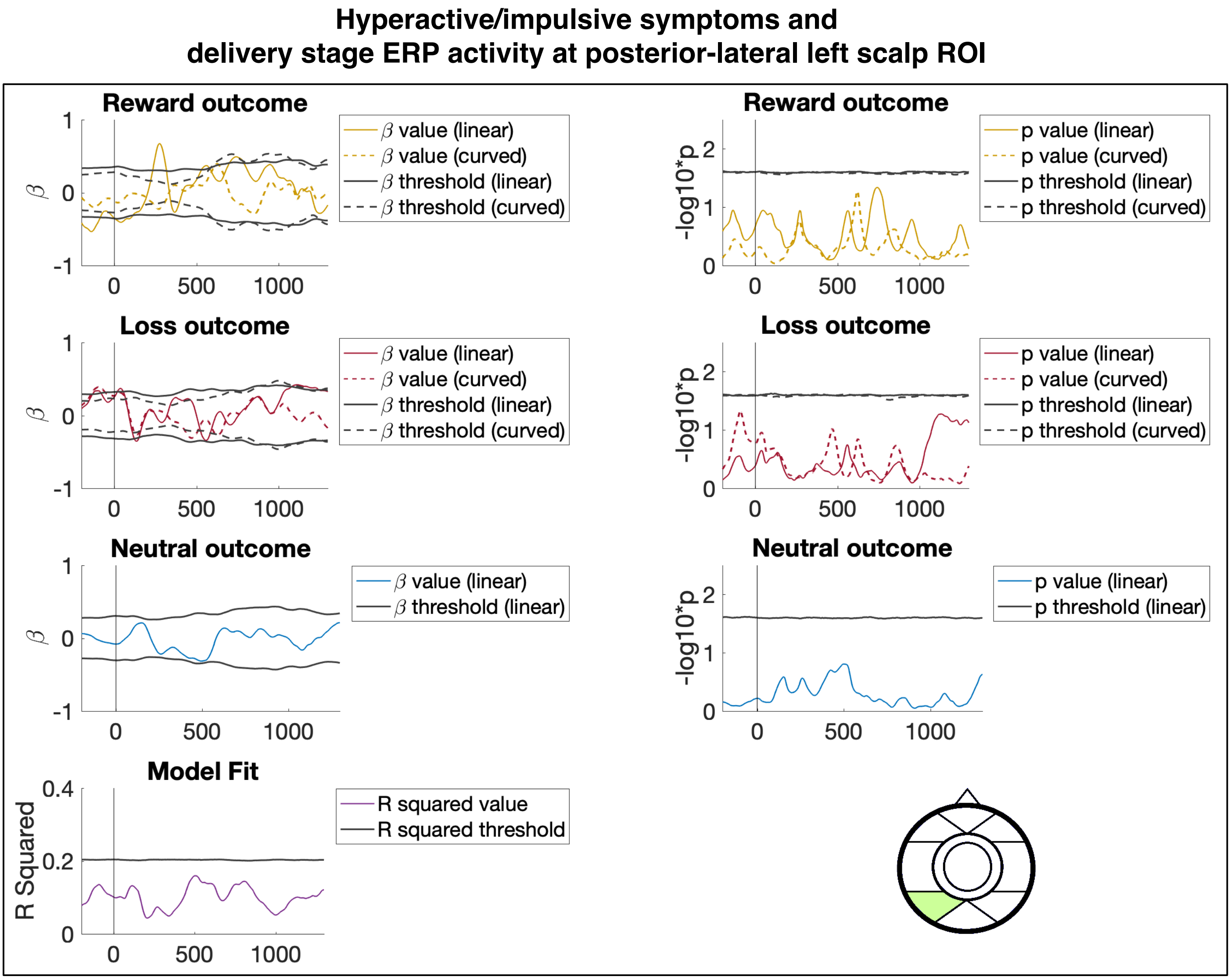


Figure S12. Association between delivery stage ERP activity and hyperactive/impulsive symptoms at the posterior-lateral left scalp ROI.

**Hyperactive/impulsive symptoms and delivery stage ERP activity at occipital-medial scalp ROI**

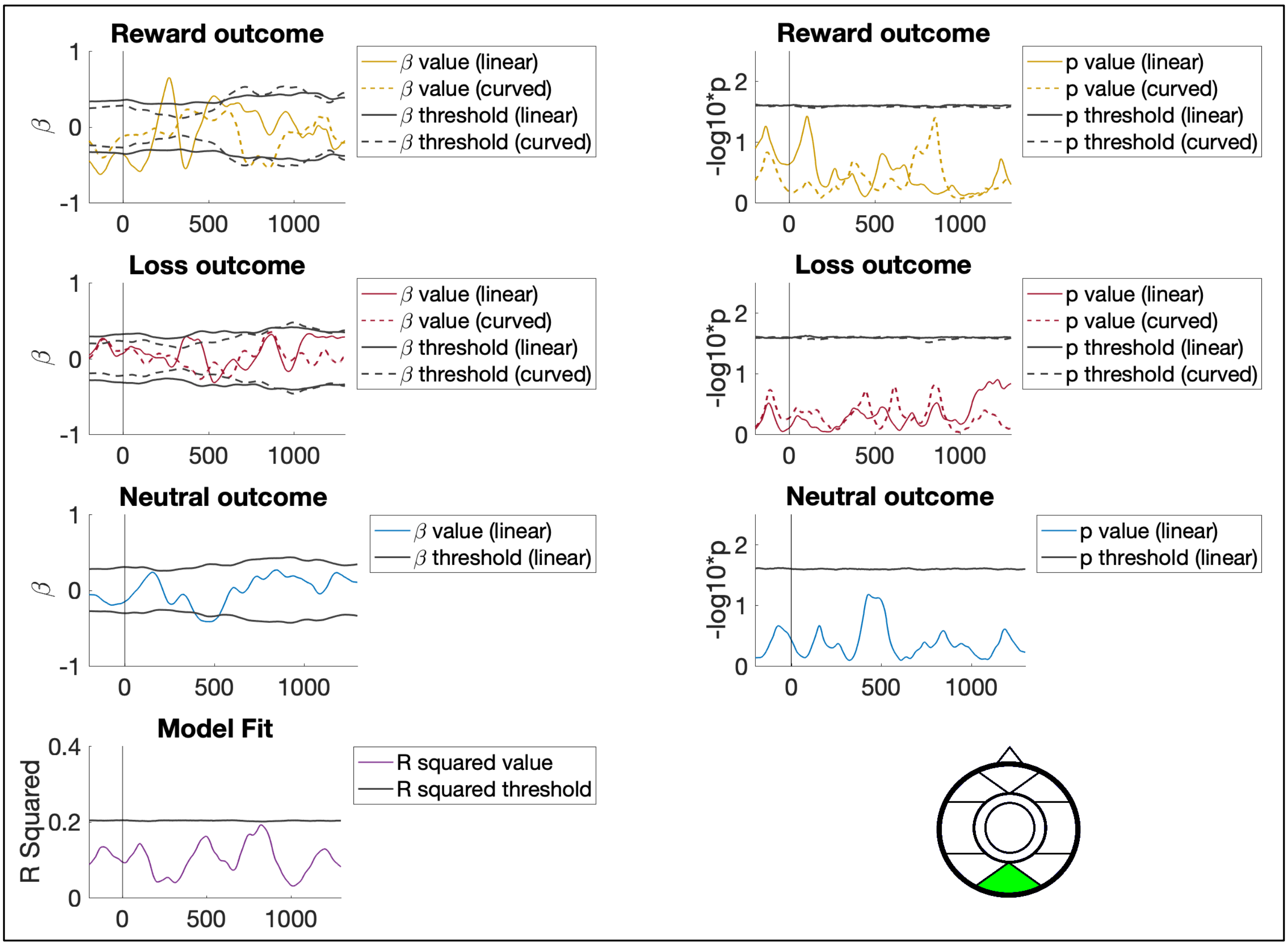


Figure S13. Association between delivery stage ERP activity and hyperactive/impulsive symptoms at the occipital-medial scalp ROI.

**Hyperactive/impulsive symptoms and delivery stage ERP activity at posterior-lateral right scalp ROI**

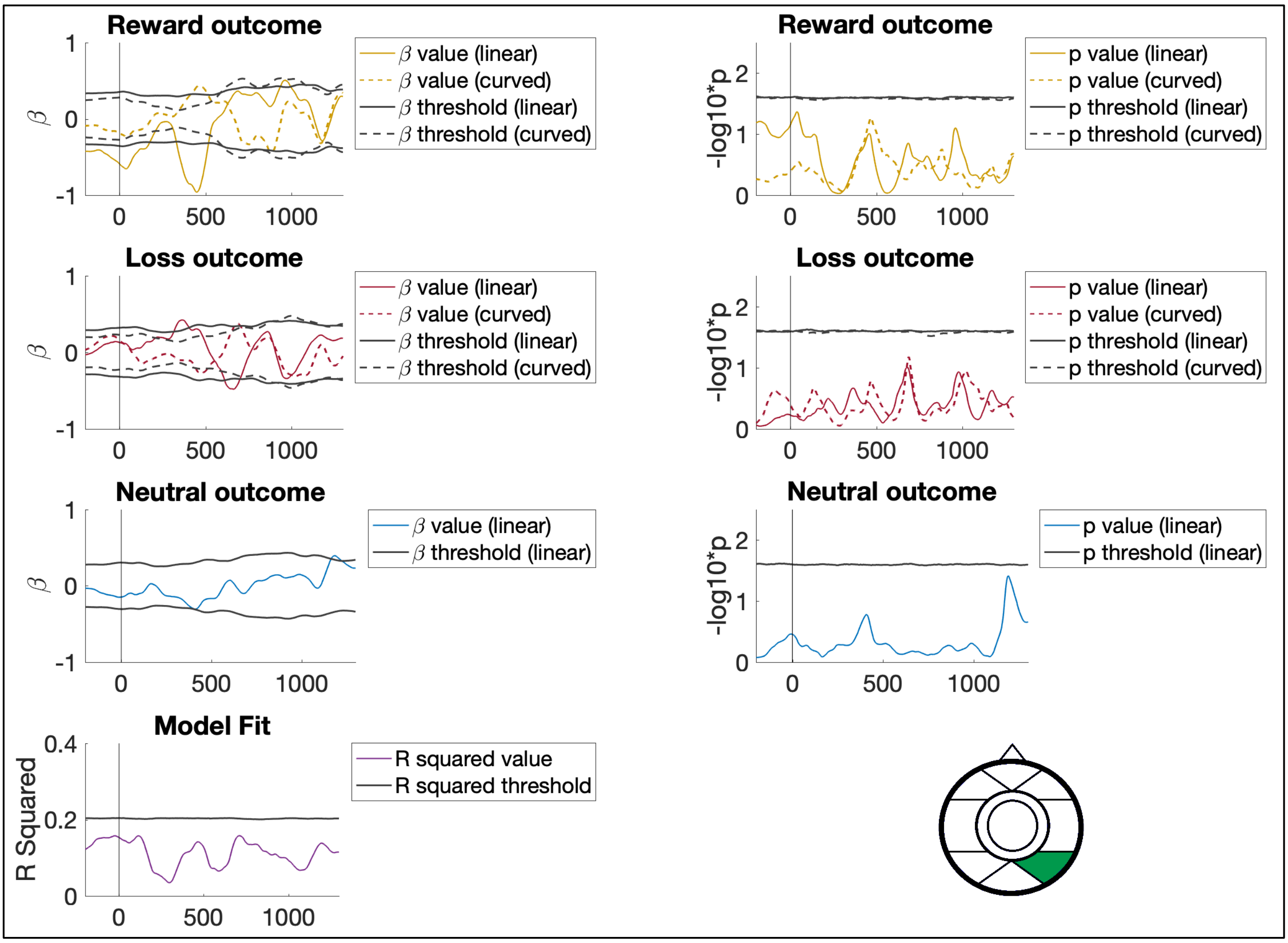


Figure S14. Association between delivery stage ERP activity and hyperactive/impulsive symptoms at the posterior-lateral right scalp ROI.

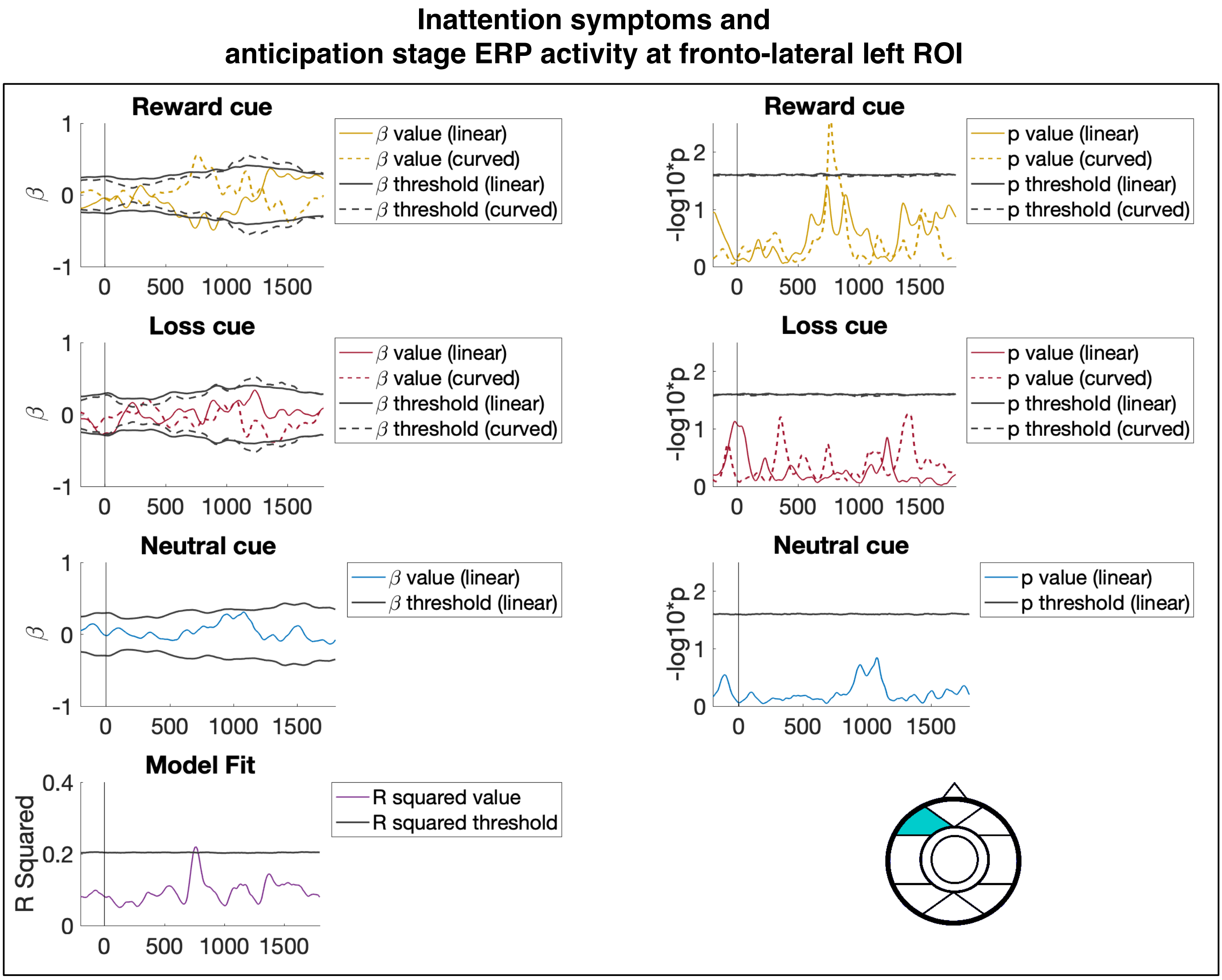


Figure S15. Association between anticipation stage ERP activity and inattention symptoms at the fronto-lateral left scalp ROI.

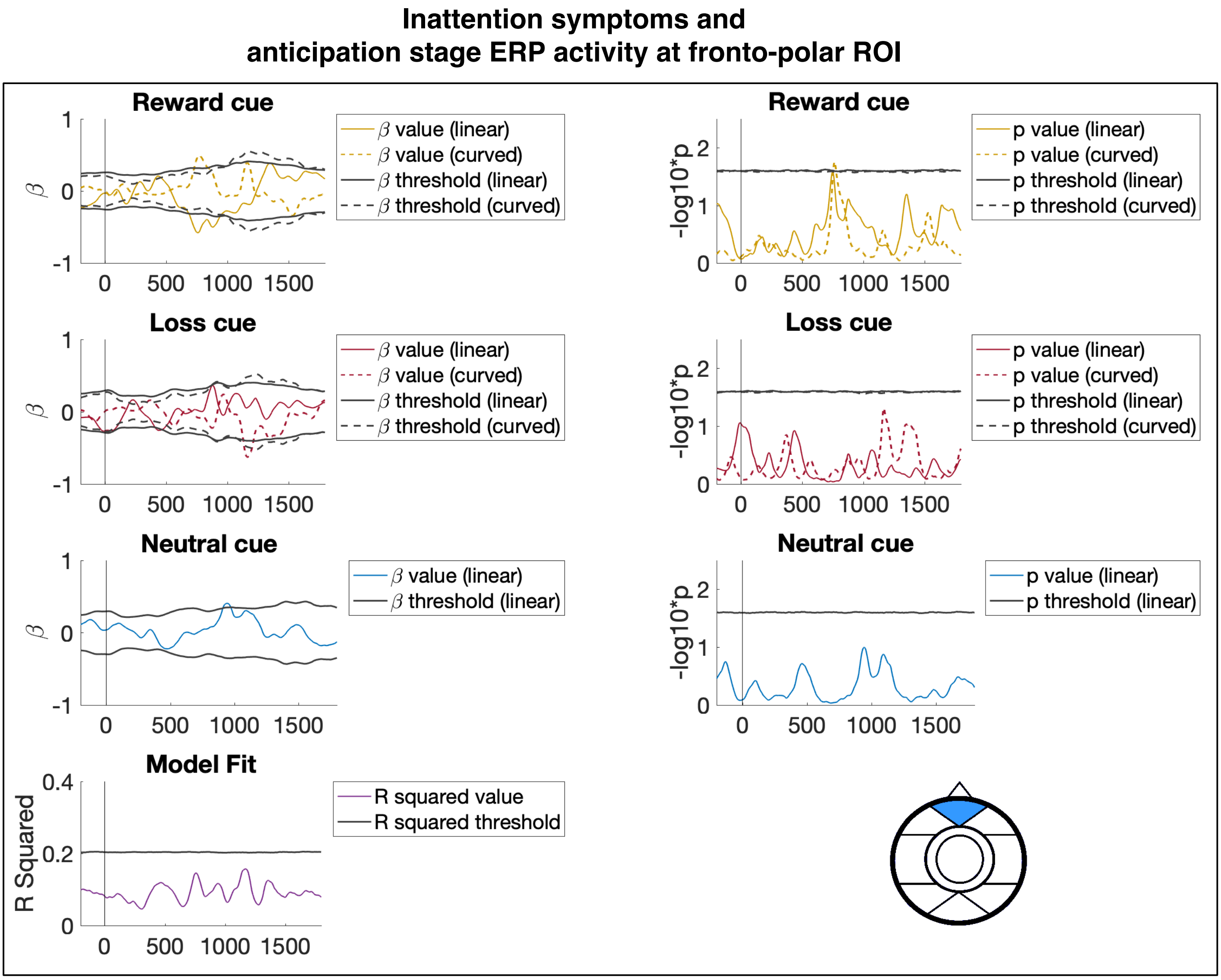


Figure S16. Association between anticipation stage ERP activity and inattention symptoms at the fronto-polar scalp ROI.

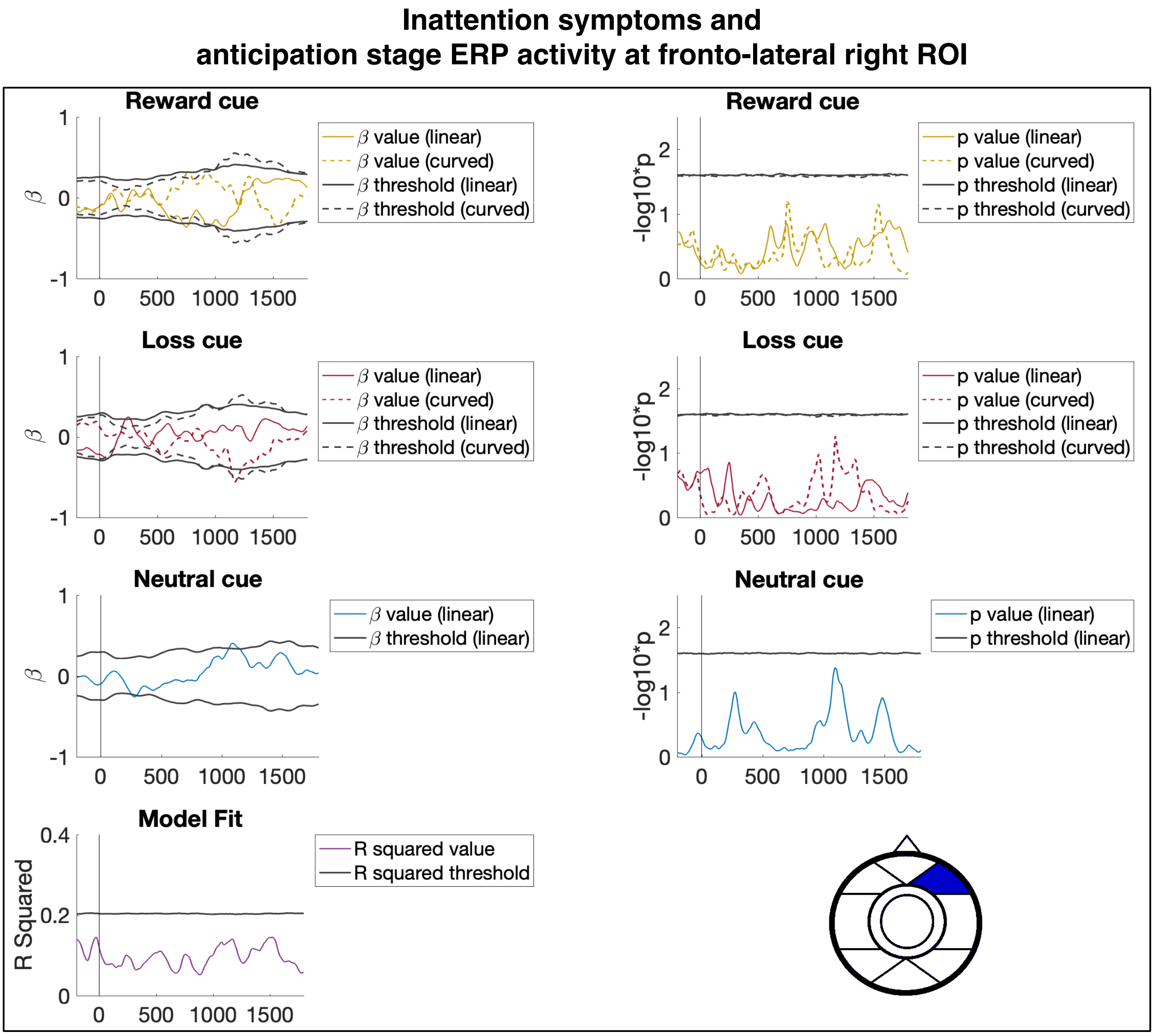


Figure S17. Association between anticipation stage ERP activity and inattention symptoms at the fronto-lateral right scalp ROI.

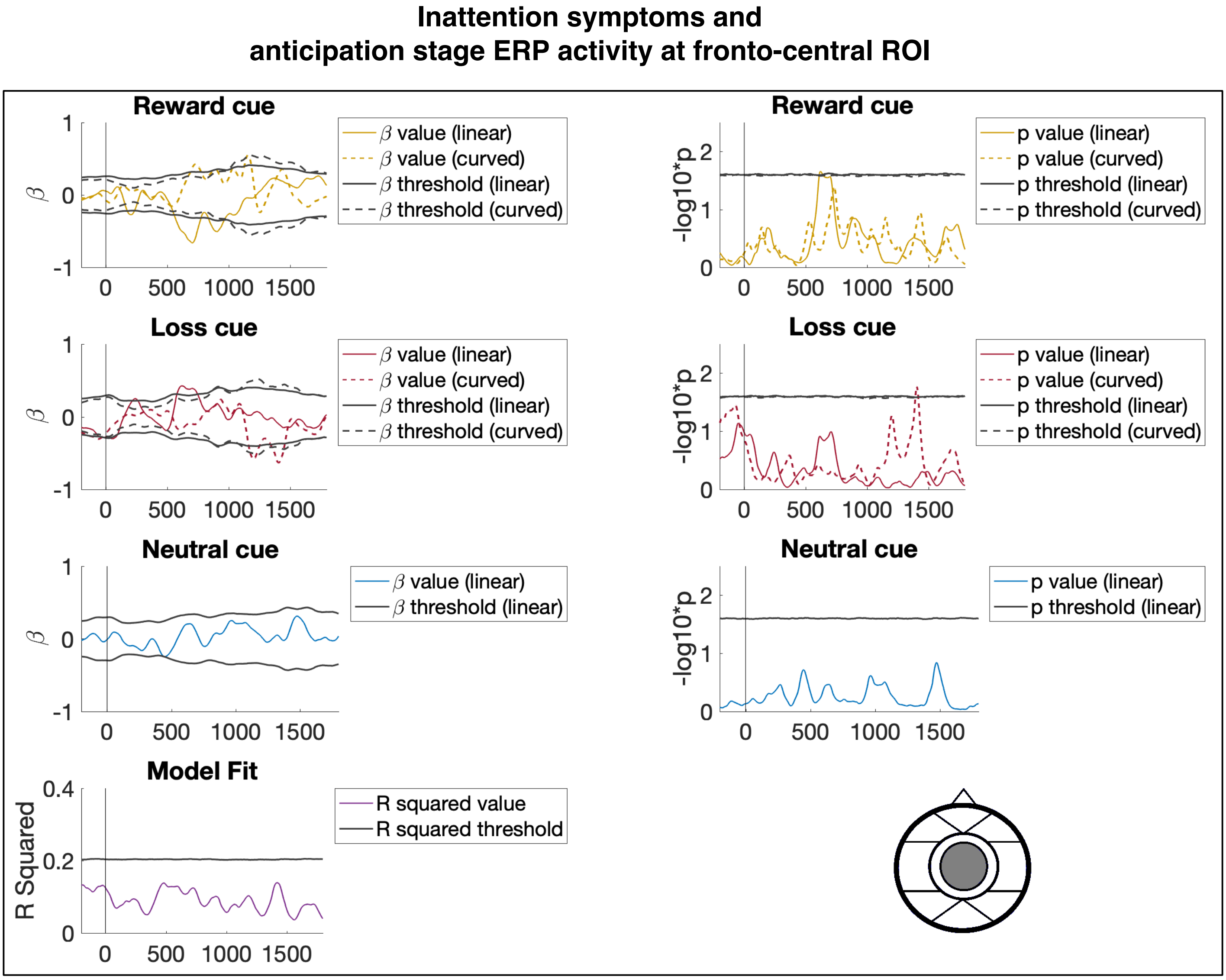


Figure S18. Association between anticipation stage ERP activity and inattention symptoms at the fronto-central scalp ROI.

**Inattention symptoms and anticipation stage ERP activity at posterior-lateral left ROI**

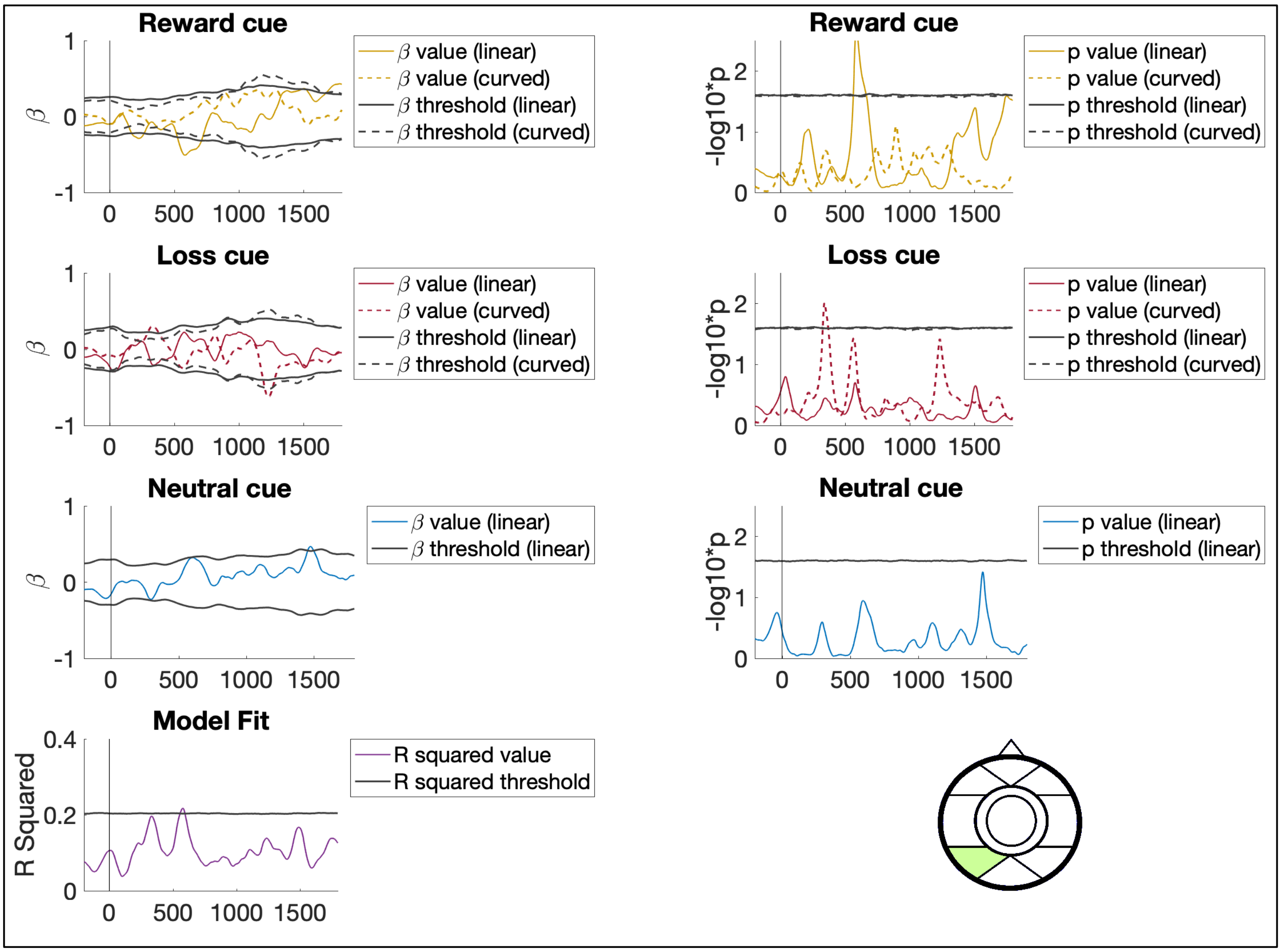


Figure S19. Association between anticipation stage ERP activity and inattention symptoms at the posterior-lateral left scalp ROI.

**Inattention symptoms and anticipation stage ERP activity at occipital-medial ROI**

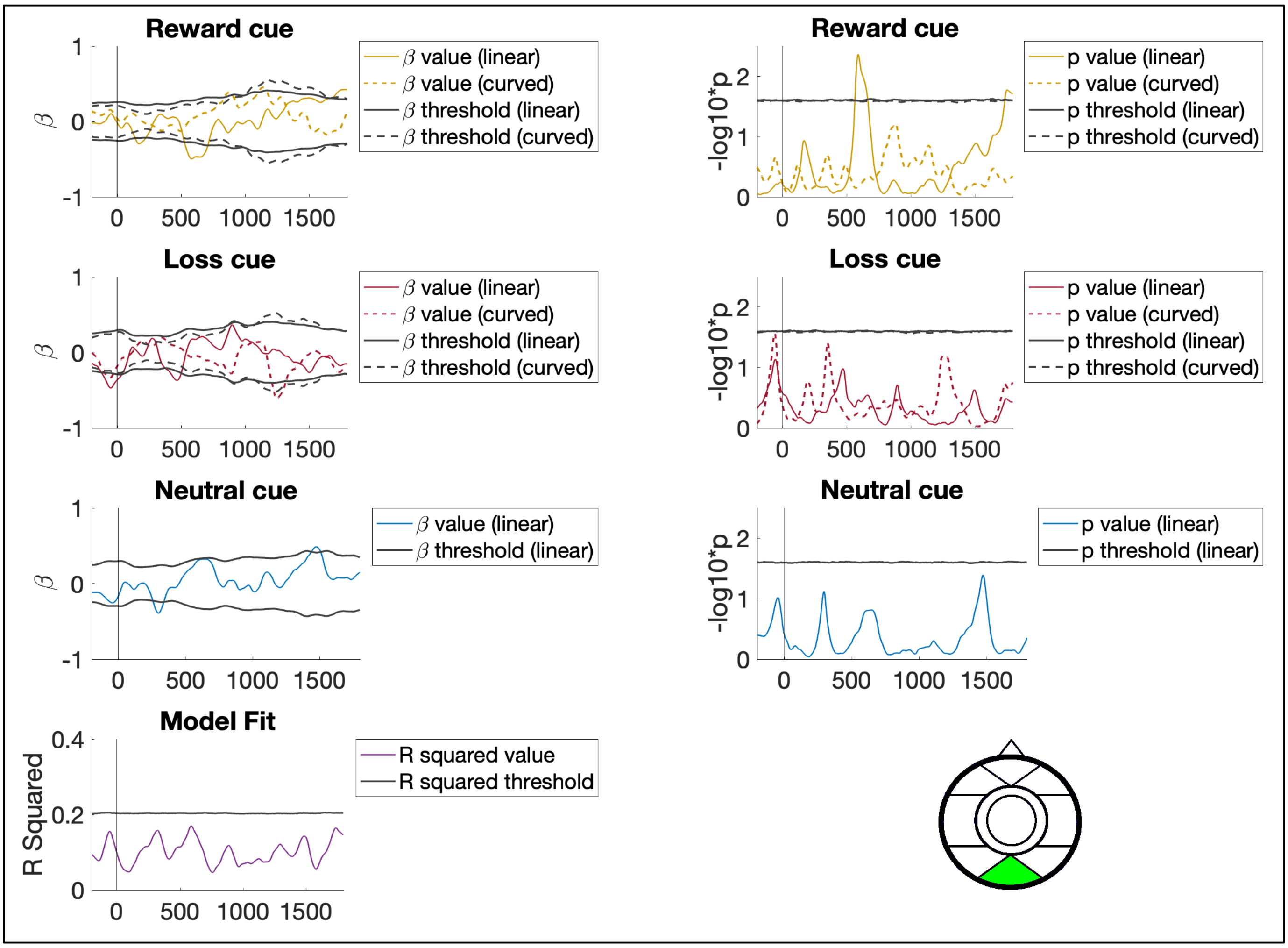


Figure S20. Association between anticipation stage ERP activity and inattention symptoms at the occipital-medial scalp ROI.

**Inattention symptoms and anticipation stage ERP activity at posterior-lateral right ROI**

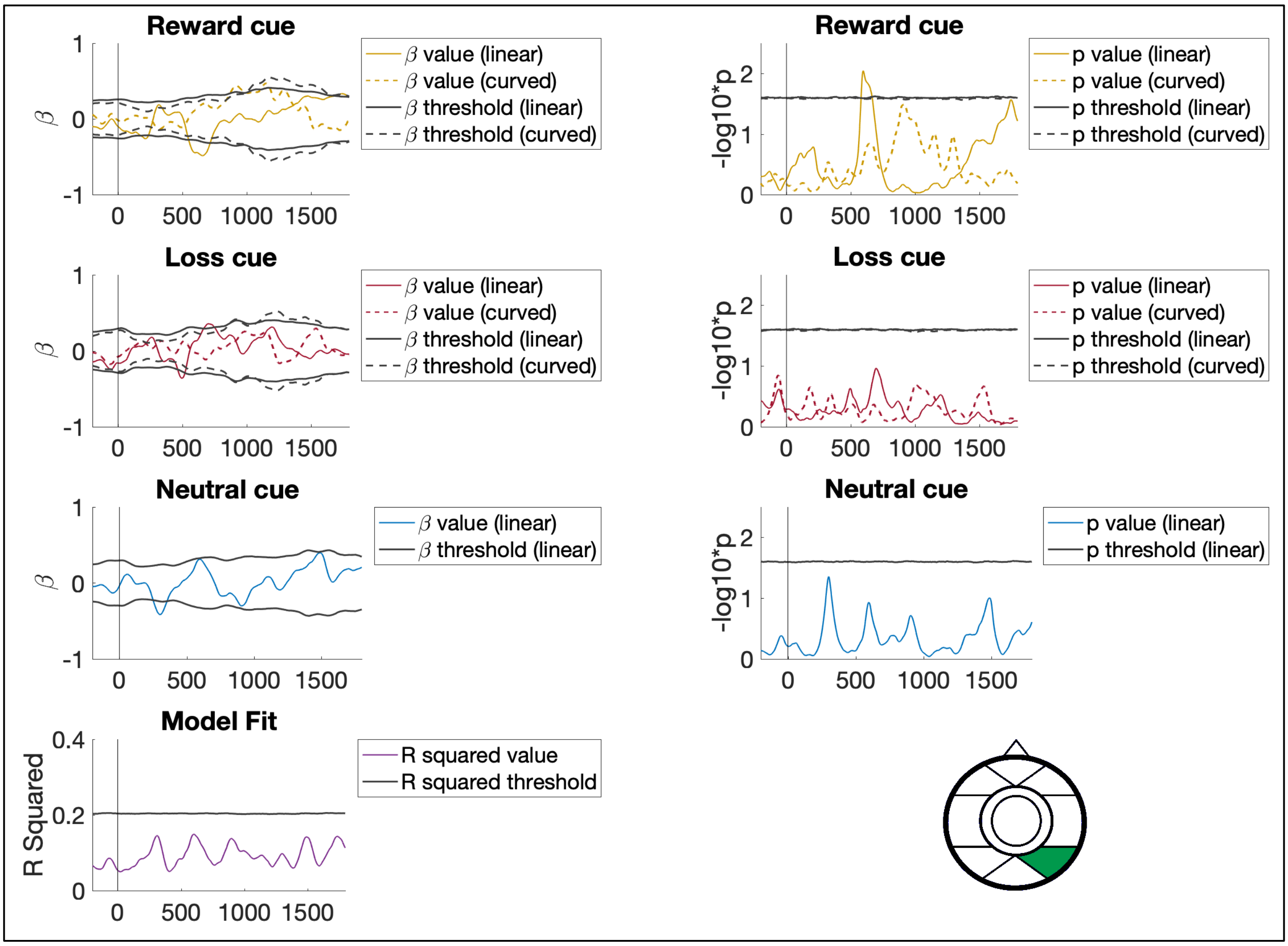


Figure S21. Association between anticipation stage ERP activity and inattention symptoms at the posterior-lateral right scalp ROI.

### Inattention symptoms and
### delivery stage ERP activity at fronto-lateral left ROI

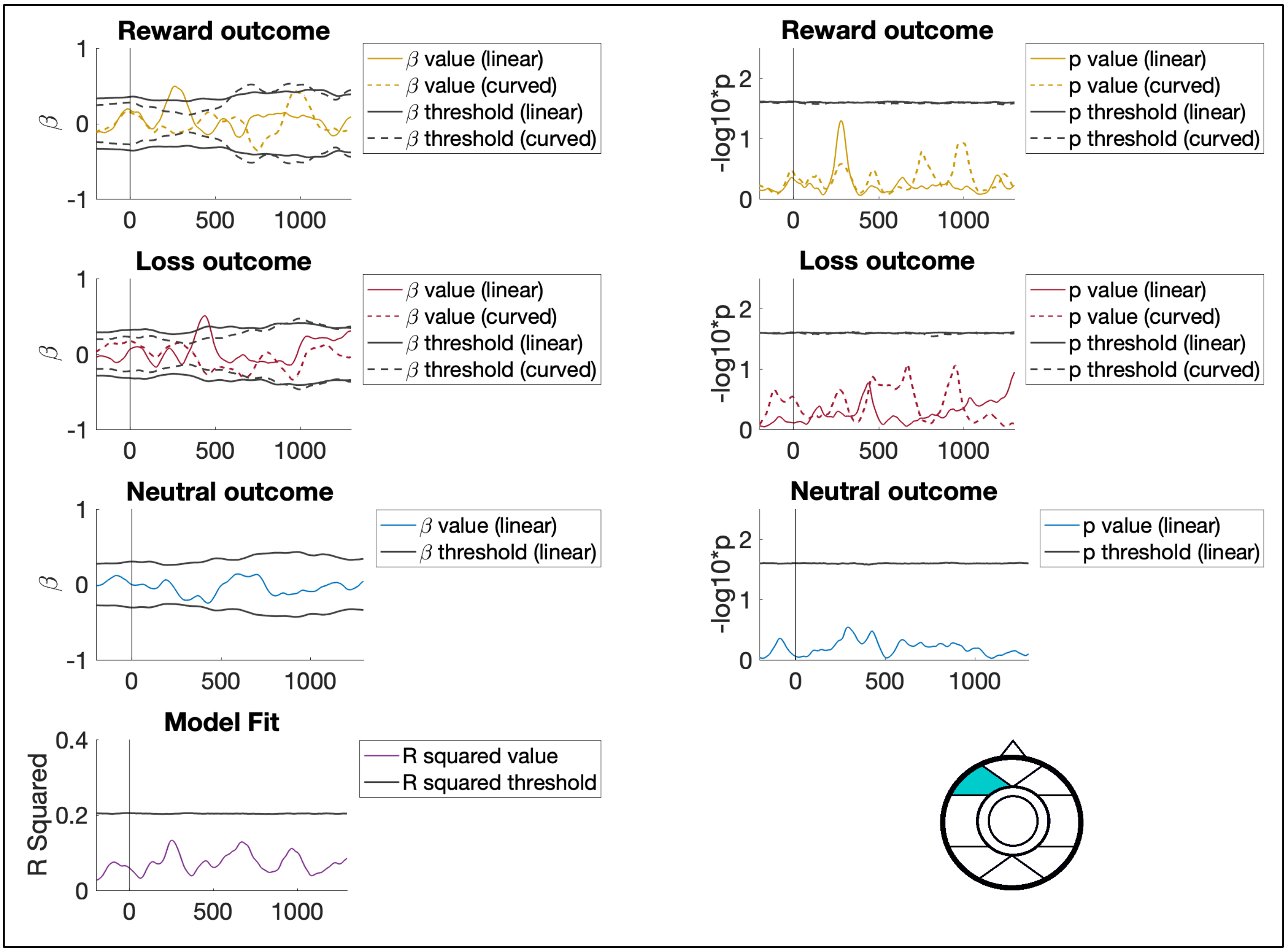


Figure S22. Association between delivery stage ERP activity and inattention symptoms at the fronto-lateral left scalp ROI.

**Inattention symptoms and delivery stage ERP activity at fronto-polar ROI**

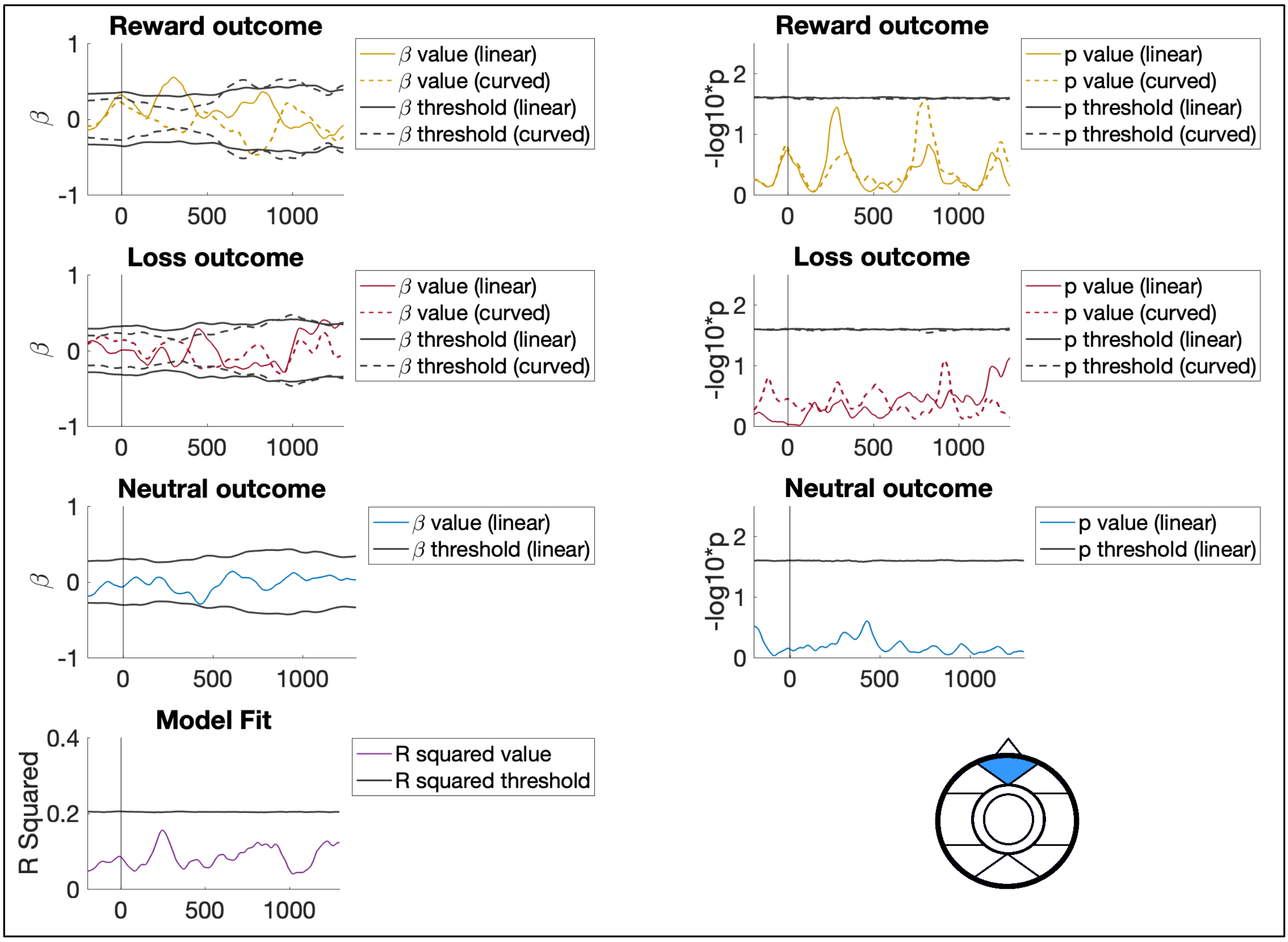


Figure S23. Association between delivery stage ERP activity and inattention symptoms at the fronto-polar scalp ROI.

**Inattention symptoms and delivery stage ERP activity at fronto-lateral right ROI**

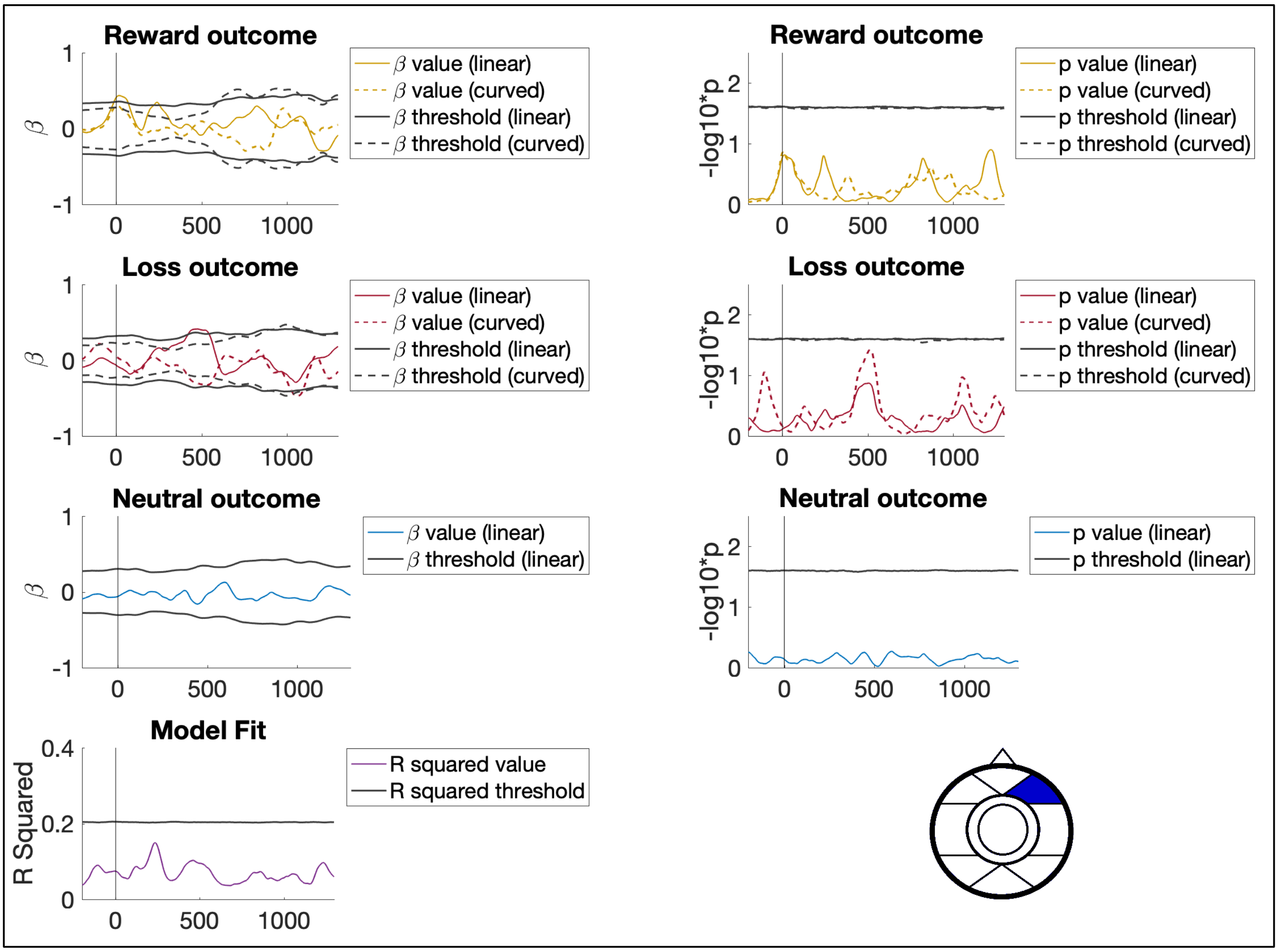


Figure S24. Association between delivery stage ERP activity and inattention symptoms at the fronto-lateral right scalp ROI.

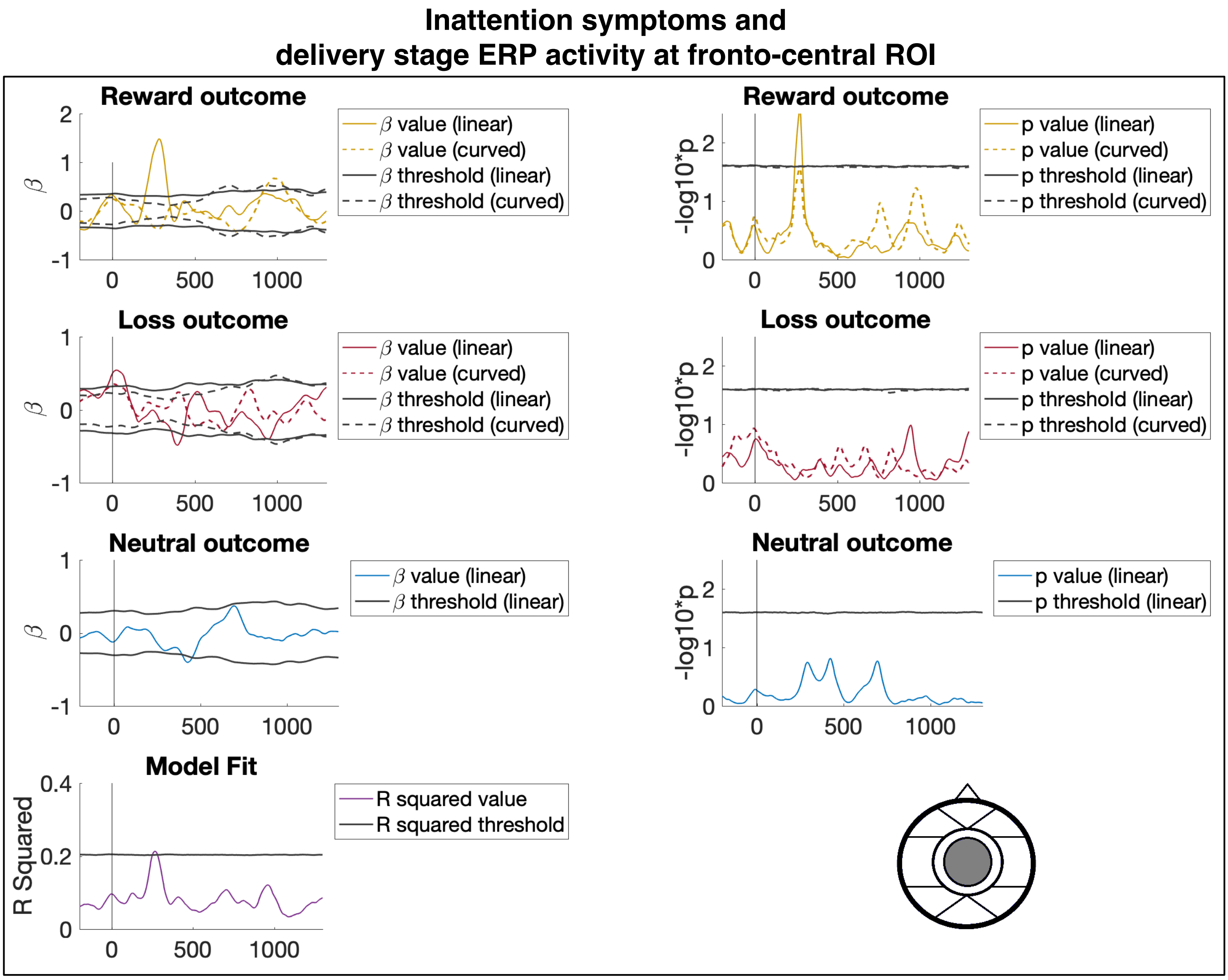


Figure S25. Association between delivery stage ERP activity and inattention symptoms at the fronto-central scalp ROI.

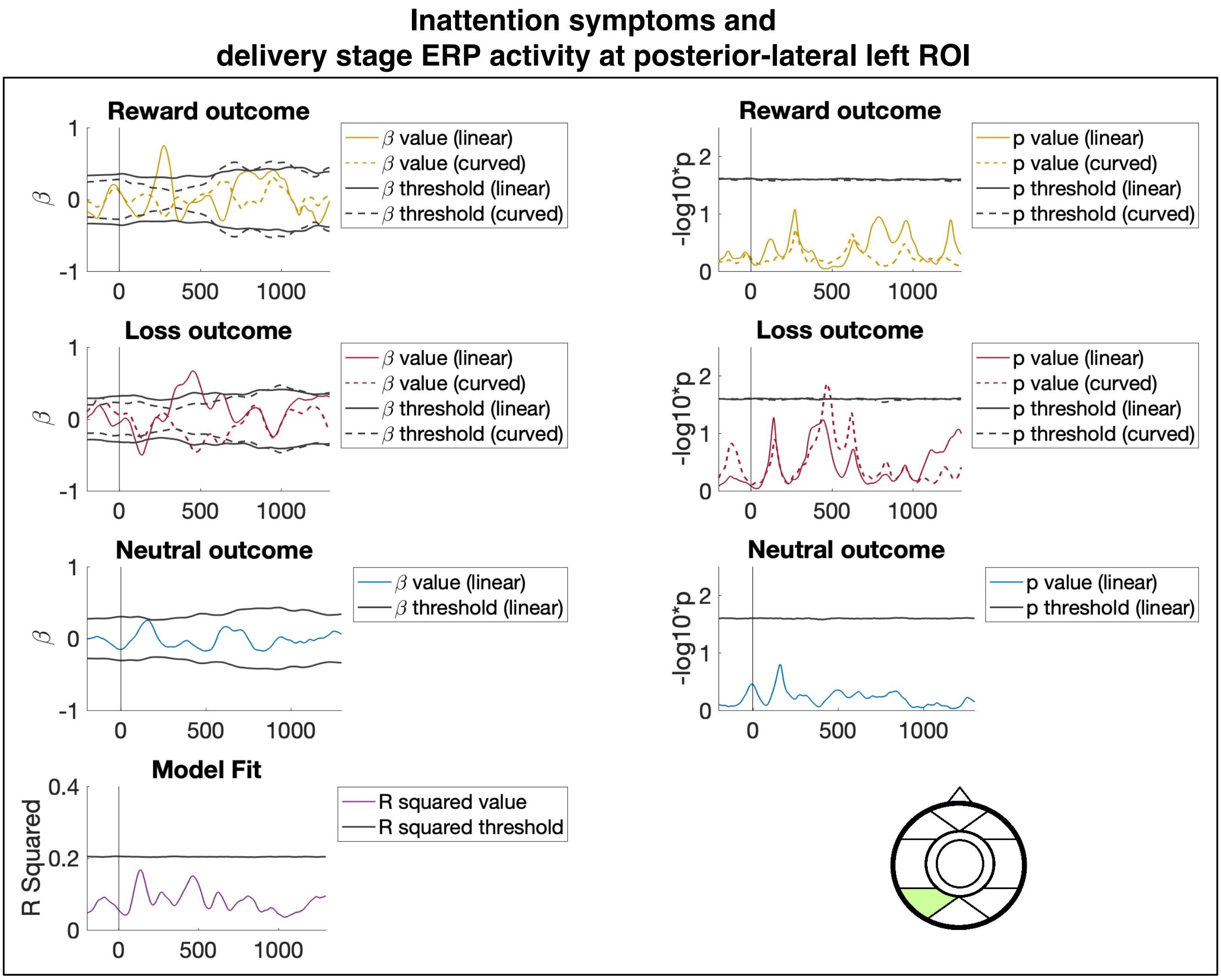


Figure S26. Association between delivery stage ERP activity and inattention symptoms at the posterior-lateral left scalp ROI.

**Inattention symptoms and delivery stage ERP activity at occipital-medial ROI**

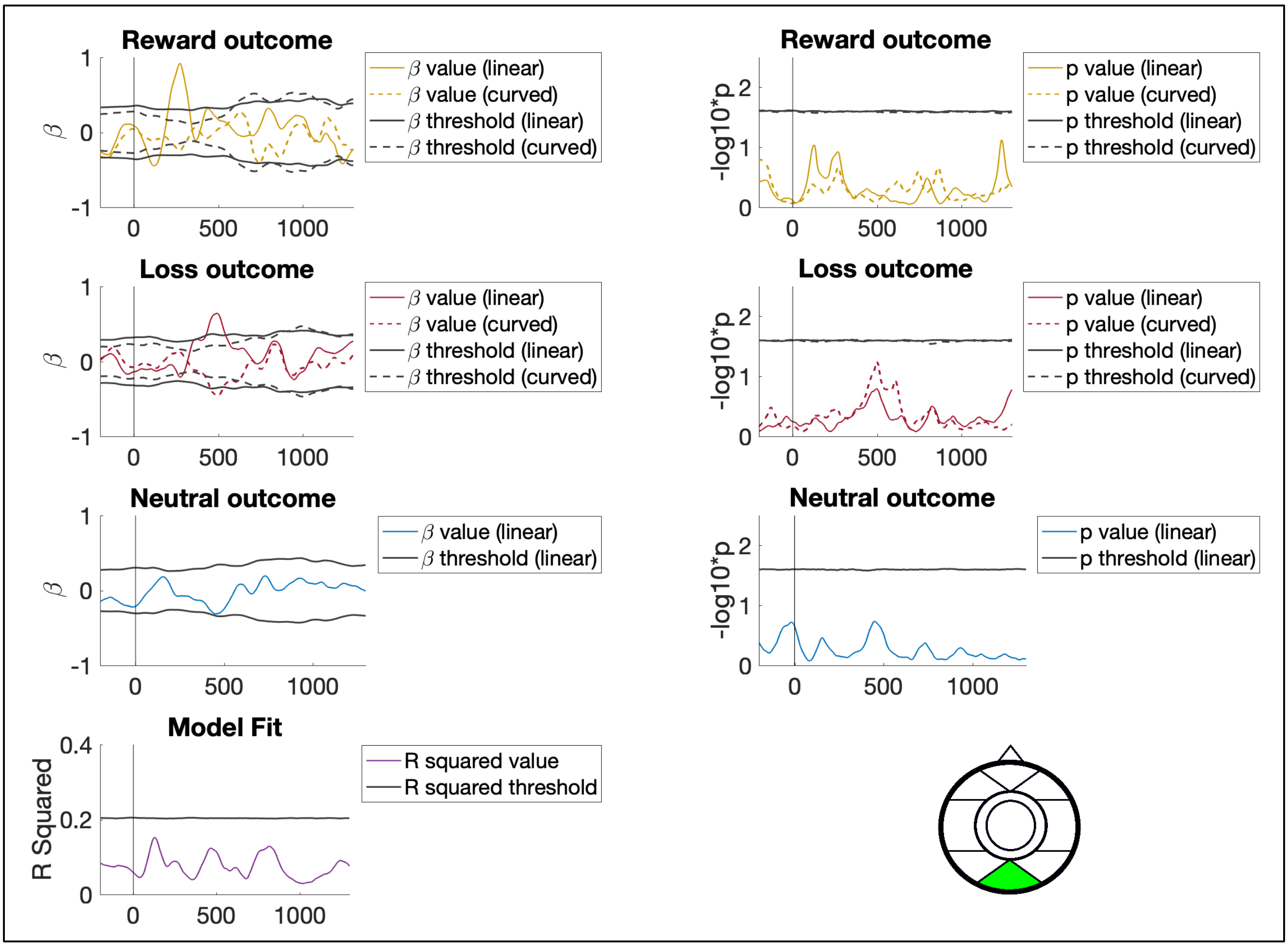


Figure S27. Association between delivery stage ERP activity and inattention symptoms at the occipital-medial left scalp ROI.

**Inattention symptoms and delivery stage ERP activity at posterior-lateral right ROI**

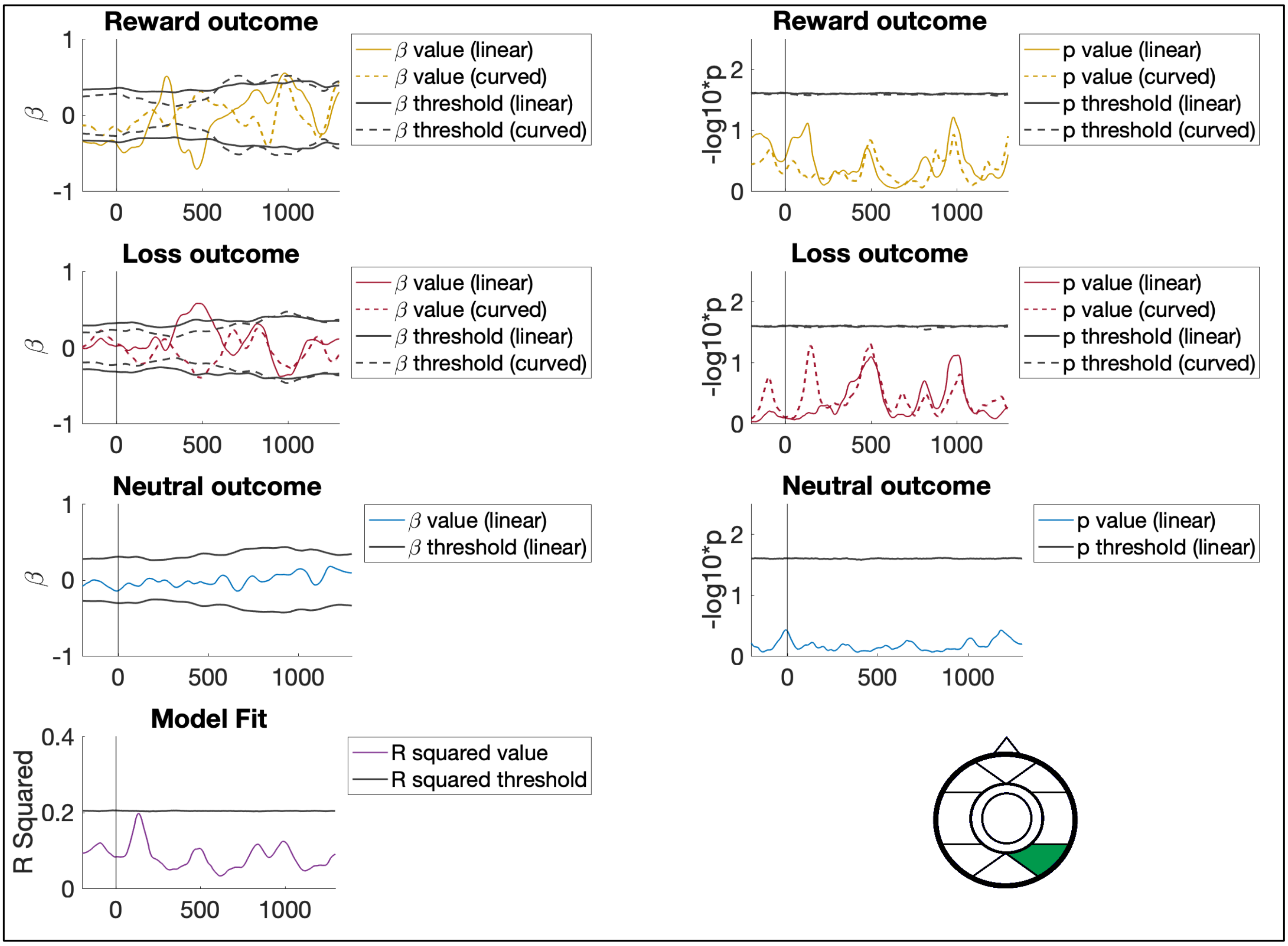


Figure S28. Association between delivery stage ERP activity and inattention symptoms at the posterior-lateral right scalp ROI.
